# Diversity and complexity of the large surface protein family in the compacted genomes of various *Pneumocystis* species

**DOI:** 10.1101/814236

**Authors:** Liang Ma, Zehua Chen, Da Wei Huang, Ousmane H. Cissé, Jamie L. Rothenburger, Alice Latinne, Lisa Bishop, Robert Blair, Jason M. Brenchley, Magali Chabé, Xilong Deng, Vanessa Hirsch, Rebekah Keesler, Geetha Kutty, Yueqin Liu, Daniel Margolis, Serge Morand, Bapi Pahar, Li Peng, Koen K.A. Van Rompay, Xiaohong Song, Jun Song, Antti Sukura, Sabrina Thapar, Honghui Wang, Christiane Weissenbacher-Lang, Jie Xu, Chao-Hung Lee, Claire Jardine, Richard A. Lempicki, Melanie T. Cushion, Christina A. Cuomo, Joseph A. Kovacs

## Abstract

*Pneumocystis,* a major opportunistic pathogen in patients with a broad range of immunodeficiencies, contains abundant surface proteins encoded by a multi-copy gene family, termed the major surface glycoprotein (Msg) gene superfamily. This superfamily has been identified in all *Pneumocystis* species characterized to date, highlighting its important role in *Pneumocystis* biology. In this report, through a comprehensive and in-depth characterization of 459 *msg* genes from 7 *Pneumocystis* species, we demonstrate, for the first time, the phylogeny and evolution of conserved domains in Msg proteins, and provide detailed description of the classification, unique characteristics and phylogenetic relatedness of five Msg families. We further describe the relative expression levels of individual *msg* families in two rodent *Pneumocystis* species, the substantial variability of the *msg* repertoires in *P. carinii* from laboratory and wild rats, and the distinct features of the expression site for the classic *msg* genes in *Pneumocystis* from 8 mammalian host species. Our analysis suggests a wide variety of functions for this superfamily, not only conferring antigenic variation to allow immune evasion but also mediating life-stage development, optimizing cell mobility and adhesion, and adapting to specific host niches or environmental conditions. This study provides a rich source of information that lays the foundation for the continued experimental exploration of the functions of the Msg superfamily in *Pneumocystis* biology.

## INTRODUCTION

*Pneumocystis* continues to be a major cause of disease in humans with immunodeficiencies, especially those with HIV/AIDS and organ transplants, and is being seen with increasing frequency in patients treated with immunodepleting monoclonal antibodies. As an atypical fungus, *Pneumocystis* has highly adapted to the mammalian lung environment (1), with a high level of host specificity; *P. jirovecii* infects humans, *P. carinii* infects Norway rats (*Rattus norvegicus*), and *P. murina* infects house mice (*Mus musculus*). In addition, *Pneumocystis* cell walls are structurally unique and differ significantly from typical fungal cell walls that are comprised of polysaccharides (mainly glucan and chitin) and highly mannosylated proteins. Both genomic and experimental analyses have shown the absence of chitin and outer chain N-mannans in *Pneumocystis* cell walls (1). Furthermore, beta-1,3-glucan is absent in the trophic form while present but masked in the cyst form of *Pneumocystis* (2).

An integral component of the *Pneumocystis* cell wall in both the cyst and trophic forms is the major surface glycoprotein (Msg) (also known as gp95, gp115, gp120 and gpA) (3-8). Ever since its identification in 1982 (9), Msg has been a focus of research, in part because it is the most abundant *Pneumocystis* protein as assessed by SDS-PAGE. Msg is present in all *Pneumocystis* species studied to date (3, 4, 6, 7, 10, 11) and appears to play an important role in pathogen-host interactions as well as evasion of host immune responses. Based on studies of *Pneumocystis* in humans, rats and mice, Msg is encoded by a multi-copy gene family with an estimated ∼30-100 copies per genome (5, 6, 8, 10, 12). *Msg* genes (up to ∼3 kb each) are closely related to but clearly distinct from each other, and are clustered together in the subtelomeric regions of multiple chromosomes (1, 13) (Supplemental Text). While there is no apparent variation in the *msg* repertoire among laboratory-bred *P. murina* or *P. carinii* isolates, extensive variation is present among *P. jirovecii* isolates (14).

Recently, we utilized long-read sequencing technology (15, 16) to identify the most complete set to date of *msg* genes in three *Pneumocystis* species (*P. jirovecii, P. carinii* and *P. murina*) as part of the *Pneumocystis* genome project (1). Based on our studies, each *Pneumocystis* genome encodes approximately 60-180 *msg* genes, depending on the species, including the classical *msg* genes, *msg*-related (termed *msr*) genes, and additional related genes. These genes are collectively termed the *msg* superfamily. We have previously reported on the first systematic classification of the *msg* superfamily (1), but did not provide a detailed description of the unique characteristics and phylogenetic relationships of individual domains and families or subfamilies. A recent report identified a small subset of *msg* genes in *P. jirovecii* from a single patient, and described potential mechanisms of recombination, but this report did not include any other *Pneumocystis* species (17).

In this report, we expanded our published analysis (1) to include *msg* genes in *Pneumocystis* from other mammalian host species. The goals of the current report were to: 1) identification of *msg* genes from *P. oryctolagi, P. macacae, P. wakefieldiae* and *P. canis*; 2) describe the characteristics and phylogenetic evolution of individual *msg* domains; 3) illustrate the characteristics, phylogenetic revolution and relative expression levels of individual *msg* families or subfamilies; 4) compare the variability of the *msg* repertoires in *P. carinii* from laboratory and wild Norway rats; and 5) characterize the variation of the expression site or upstream conserved sequence (UCS) of the classic *msg* genes in *Pneumocystis* from 11 mammalian host species.

## RESULTS

### Sources of *Pneumocystis msg* sequences

*msg* sequences for *P. murina, P. carinii* and *P. jirovecii* were obtained primarily from our previous *msg* and genome sequencing studies (1, 15, 18). Additional *msg* sequences were identified by Sanger sequencing of cloned *msg* amplicons and next-generation sequencing of whole genomes of the following *Pneumocystis* species: *P. wakefieldiae* infecting laboratory rats, *P. carinii* infecting wild rats, *P. macacae* infecting rhesus macaques, *P. canis* infecting dogs, and *P. oryctolagi* infecting rabbits (Table 1 and Table S1). Due to the low-throughput nature and high cost of Sanger sequencing of cloned *msg* amplicons and the difficulty in assembling short reads from Illumina sequencing, only a small number of full-length *msg* genes were obtained from these species (1-13 genes per species). As whole genome assembly of these species is still in progress, the *msg* genes reported for these species are only representative, not all-inclusive. All *msg* sequences are available from Supplemental Data Sets 1 to 8 as well as the BioProject database with accession no. PRJNA560924.

**Table 1.** Summary of the *msg* superfamily members identified in *Pneumocystis* species.

| Family | No. of *msg* genes<sup>a</sup> |  |  |  |  |  |  | Characteristics<sup>b</sup> |  |  |  |  |
| --- | --- | --- | --- | --- | --- | --- | --- | --- | --- | --- | --- | --- |
|  | *P. murina* | *P. carinii* | *P. jirovecii* | *P. wakefieldiae* | *P. macacae* | *P. canis* | *P. oryctolagi* | Average gene (kb)/protein (kDa) size | No. of introns | 5'-end leader sequence | No. of domains | Expression mode<sup>c</sup> |
| *msg*-A1<sup>c</sup> | 22 | 65 | 86 | 2 | 2 | 9 | 7 | 3.2/120 | 0 | CRJE<sup>d</sup> | 9 | Mutually exclusive |
| *msg*-A2<sup>c</sup> | 14 | 53 | 0 | 6 | 0 | 0 | 0 | 2.9/218 | 1 | Highly conserved | 7-9 | Independently |
| *msg*-A3<sup>c</sup> | 6 | 3 | 33 | 2 | 3 | 1 | 0 | 3.1/117 | 1-2 | Highly variable | 9 | Independently |
| *msg*-B | 0 | 0 | 21 | 0 | 2 | 0 | 1 | 1.4/55 | 1-2 | Highly variable | 2-3 | Independently |
| *msg*-C | 6 | 1 | 2 | 2 | 1 | 0 | 0 | 1.7/60 | 1 | Highly conserved | 3 | Independently |
| *msg*-D | 1 | 1 | 20 | 1 | 4 | 2 | 1 | 2.9/111 | 1 | Highly variable | 3-6 | Independently |
| *msg*-E | 7 | 5 | 5 | 7 | 5 | 2 | 2 | 1.2/49 | 1 | Highly variable | 1 | Independently |
| Unclassified | 8 | 13 | 12 | 0 | 0 | 0 | 0 | 0.9/37 | 0-8 | Highly variable | 0-1 | Unknown |
| *Total* | 64 | 141 | 179 | 20 | 17 | 14 | 11 |  |  |  |  |  |

<sup>a</sup>Results represent a potentially complete set for *P. murina*, *P. carinii* and *P. jirovecii* as described in reference (1) but only subset
members for the other four species listed, whose genome sequencing is still in progress.
<sup>b</sup>Based on *P. murina*, *P. carinii* and *P. jirovecii* genes with complete or nearly complete reading frames.
<sup>c</sup>Subfamilies belonging to *msg*-A family.
<sup>d</sup>Conserved recombination joint element, which is unique, highly conserved among all *msg*-A1 genes.

### Characteristics and phylogenetic relationships of individual Msg domains

We identified a total of 9 conserved domains (Fig. 1 and Text S1). Classic Msg (Msg-A1) proteins contain 5 domains that presumably arose as gene duplication. Based on phylogenetic trees constructed using only these 5 domains, we found that each forms its own cluster, regardless of the origin of the species of the domains (Fig. S1). This strongly suggests that the most recent common ancestor to these *Pneumocystis* species already had developed this Msg domain structure and organization, and that subsequently these domains evolved with no further duplication or recombination among domains across or within species. We also found that within each of these 5 Msg domains, individual domains seem to cluster by *Pneumocystis* species, suggesting that significant Msg family expansion occurred after the separation of *Pneumocystis* species. In addition, *P. carinii* and *P. murina* form two separate clusters with each cluster containing both species, suggesting that those two clusters arose before separation of these two species.

**FIG 1.**
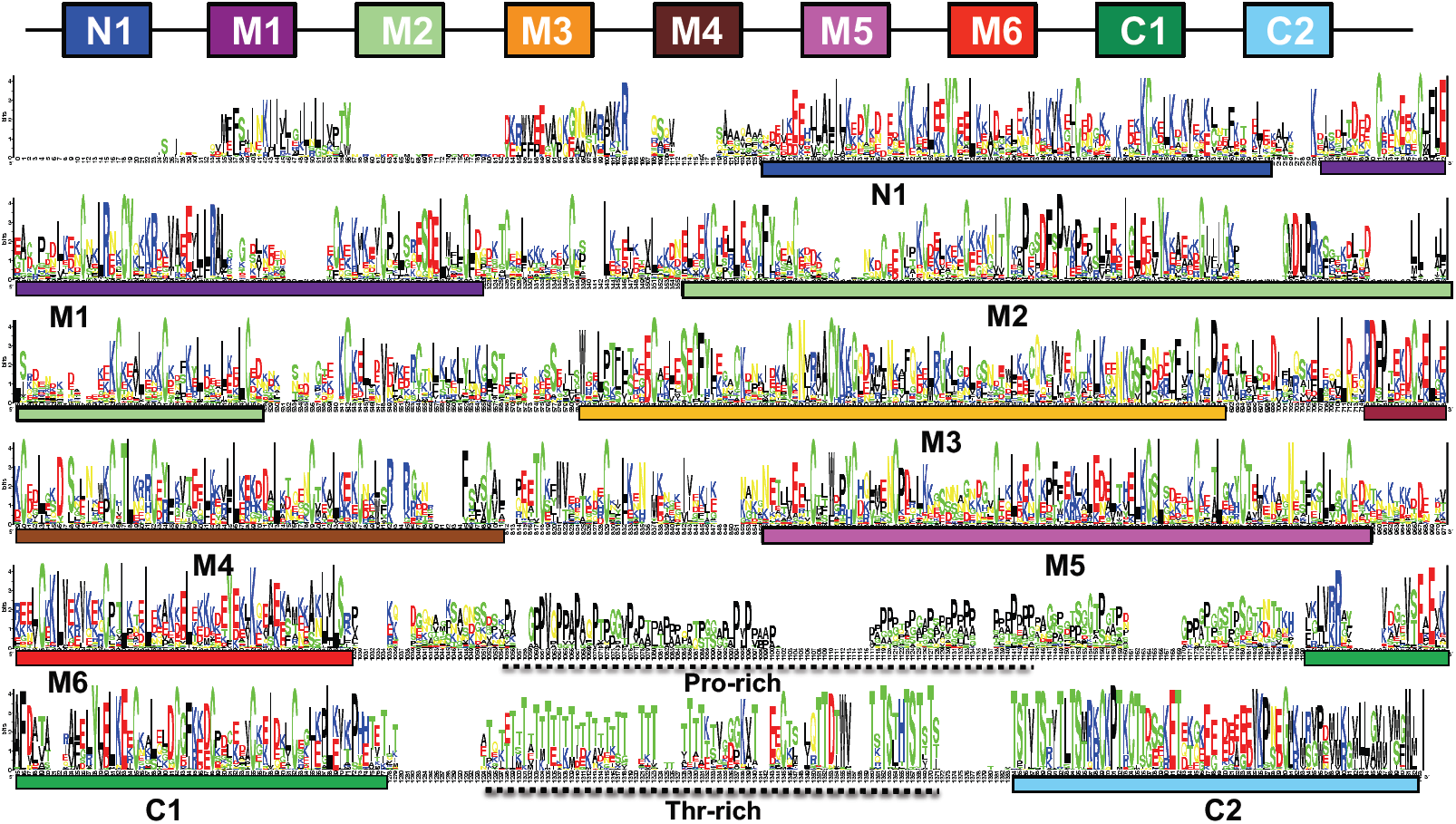
Sequence logo showing the alignment of full-length Msg proteins in *P. murina, P. carinii* and *P. jirovecii*. The known Pfam domains M1 to M5 (Pfam MSG) and C1 (extended from Pfam Msg2_C to cover a longer conserved region), three new domains (N1, M6 and C2), and Pro- and Thr-rich regions are indicated. The horizontal axis represents the position of the amino acids. The vertical axis indicates conservation of each position as measured by information content (bits). This logo is adapted from Figure 3 of reference (1) in which individual domains were shown separately without aligning with full-length proteins.

The 31 N-linked glycans from *P. carinii* Msg proteins previously identified by liquid chromatography-tandem mass spectrometry (1), mapped to 4 domains, most commonly domains M4 and M5 (each with 13 glycans), less commonly with domains M2 (2 glycans) and M3 (3 glycans).

### Unique characteristics of each Msg family and subfamily

Based on domain structure, phylogeny analysis and expression control mechanisms of the *msg* superfamily, we have previously proposed a classification of five families, named as Msg-A, Msg-B, Msg-C, Msg-D, and Msg-E (1), as summarized in Table 1. According to the chromosomal-level assemblies of the *P. murina, P. carinii* and *P. jirovecii* genomes, *msg* genes are located almost exclusively in subtelomeric regions and usually present in clusters (Text S1). Different *msg* families differ in the number of members, distribution among different *Pneumocystis* species, sequence structure (gene length, location and number of introns, and number of conserved domains), and expression control mechanisms, as summarized in Table 1. In addition, there is a bias of amino acid distribution among different Msg families (Fig. S10 and Text S1).

### Msg-A family

Msg-A family is by far the largest among the 5 families of the Msg superfamily. This family is divided into three subfamilies of Msg-A1, Msg-A2 and Msg-A3 (Fig. 2 and Figs. S2-S4). In addition to difference in phylogenetic relationships, these three subfamilies have significant differences in the expression control mechanism and sequence structure of the 5’-end leader.

**FIG 2.**
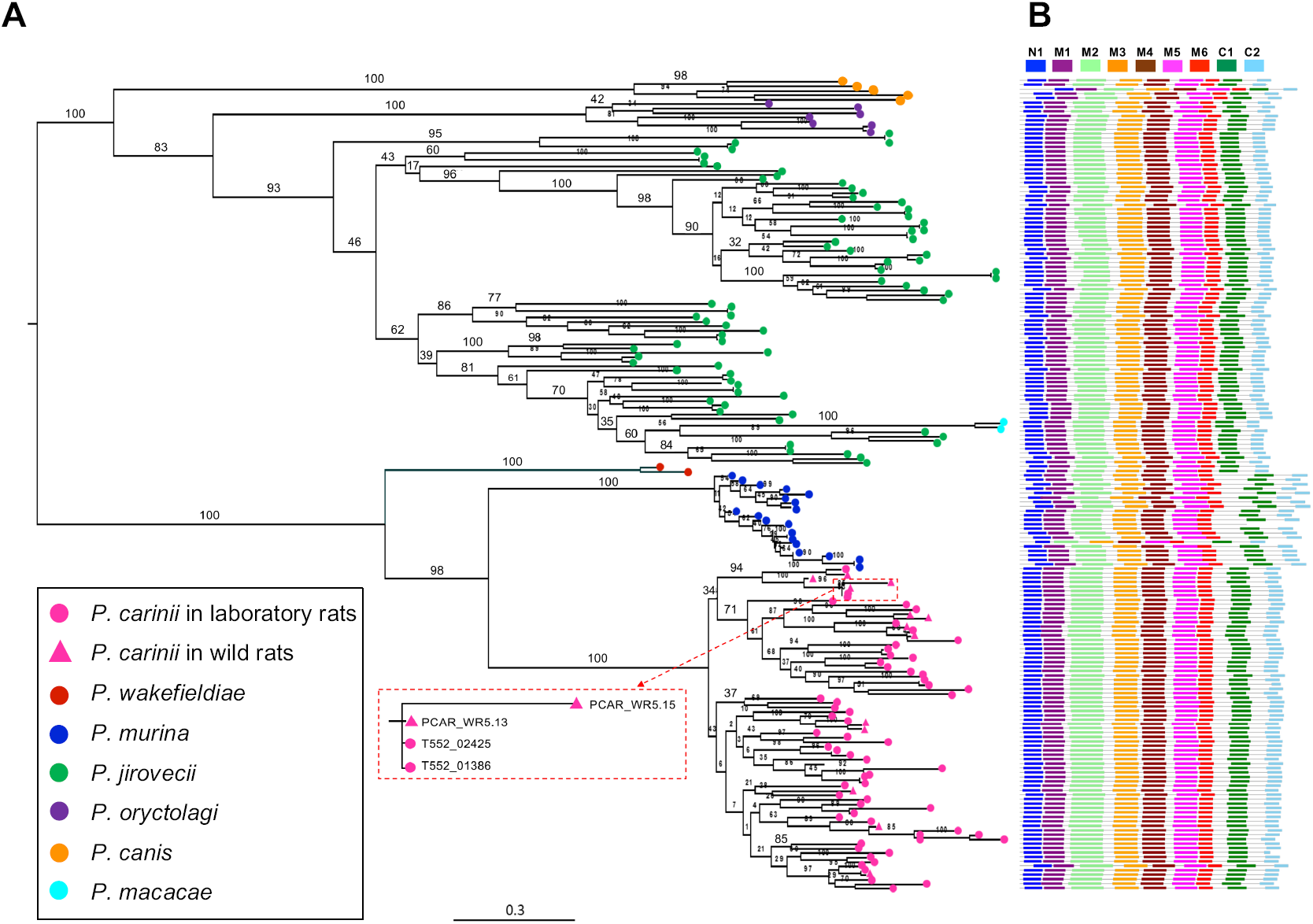
Phylogenetic tree and conserved domain structure of classic Msg genes (Msg-A1 subfamily). (**A**) A maximum-likelihood (ML) tree constructed using deduced full-length protein sequences of *msg*-A1 genes. Different *Pneumocystis* species are color-coded as indicated at the bottom left corner. *P. carinii* from laboratory rats and wild rats are indicated by pink dots and pink triangles, respectively. Only one of the 13 Msg proteins from the wild rat (PCAR_WR5.13) was nearly identical (one amino acid difference) to two Msg proteins *in P. carinii* from laboratory rats (T552_02425 and T552_01386) as shown in the red box with dashed-line. Numbers on the branch nodes indicate bootstrap support values. All sequences shown are available from Supplemental Data Set 1. (**B**) Schematic representations of conserved Msg domains. Different domains are color-coded as indicated at the top. Each row corresponds to the domain structure of the corresponding protein in panel A.

The Msg-A1 subfamily includes all classic *msg* genes. The most striking characteristics of this subfamily are its dominance among all Msg families/subfamilies across all *Pneumocystis* species (Table 1) and its unique expression control mechanism. It has been well established that expression of this subfamily is controlled by a dedicated, single-copy subtelomeric expression site, also known as the upstream conserved sequence or UCS (19-22) (Fig. 3). The UCS is expressed in fusion with a *msg* gene; the region between UCS and its downstream *msg* gene is termed the conserved recombination joint element (CRJE), which is highly conserved among all *msg-*A1 genes, and potentially serves as an anchor for recombination (23). Available data suggest that these *msg* genes are not expressed unless they are translocated downstream of and in-frame with the UCS (19-22). This mechanism allows only one *msg*-A1 gene to be expressed in a single organism at a given time, although multiple *msg-*A1 genes are expressed at the population level in immunosuppressed hosts. In phylogenetic analysis of 183 full-length *msg*-A1 genes from 7 *Pneumocystis* species (Fig. 2), these genes cluster by species; as expected, genes from all three rodent *Pneumocystis* species form a strong monophyletic group (Fig. 2). As previously noted (14), the Msg-A1 family in *P. jirovecii* is composed of two phylogenetically distinct groups; such separation is also seen in *P. murina* and *P. carinii*.

**FIG 3.**
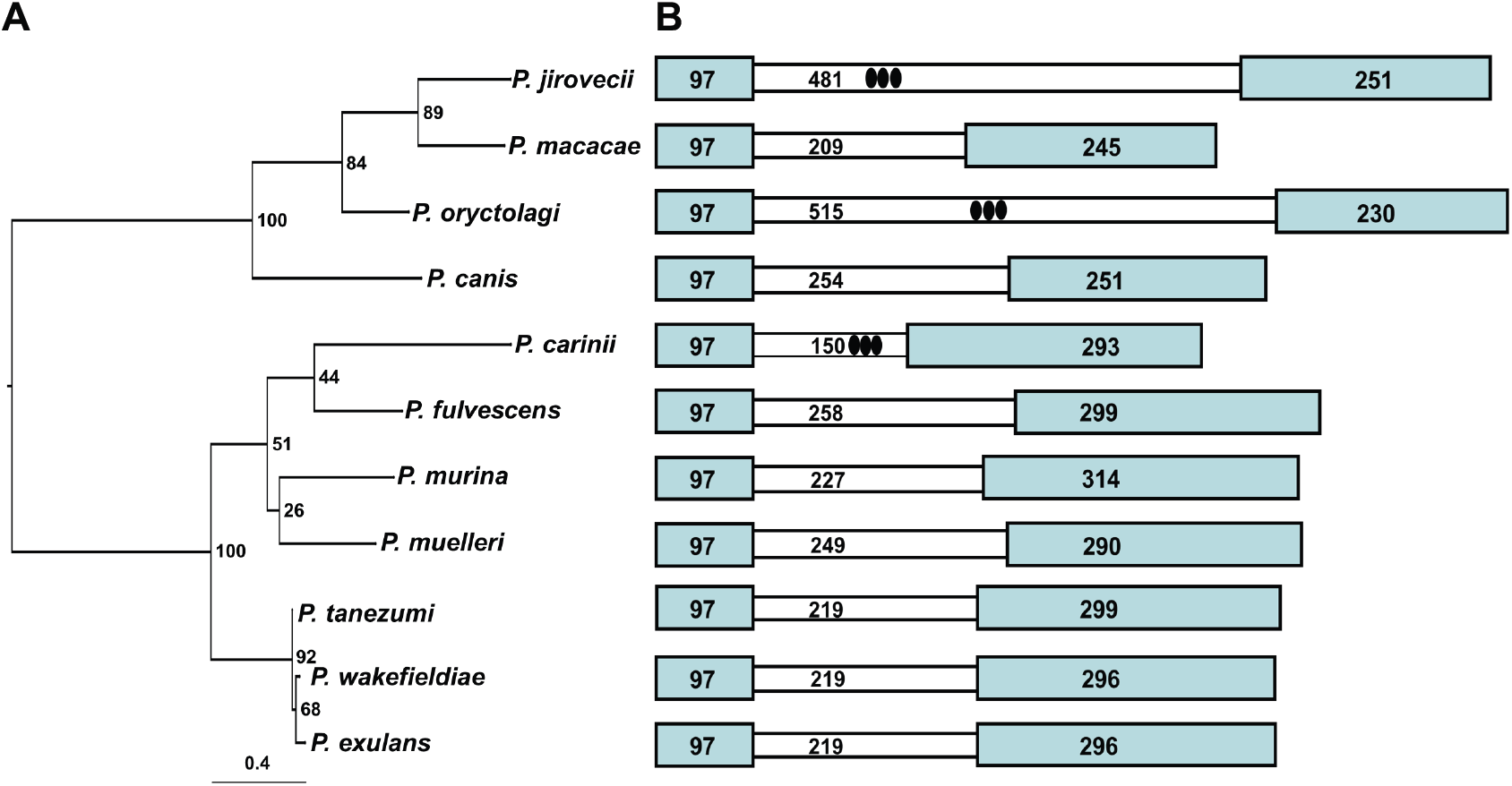
Phylogeny and sequence structure of the expression sites or upstream conserved sequences (UCSs) of *msg*-A1 genes in *Pneumocystis* from different mammalian species. (**A**) Phylogenetic relationship based on protein sequences of UCSs. Numbers on the branches indicate bootstrap support values. (**B**) Schematic representation of the UCS sequence structure. The number in each box is the sequence length (bp) for each region. The approximate location of the tandem repeats in *P. jirovecii, P. oryctolagi* and *P. carinii* are indicated by ovals, with more details on tandem repeat variation in *P. oryctolagi* and *P. carinii* shown in Fig. 6 and Fig. S13, respectively. Details about the nomenclature of *Pneumocystis* and its host species are available in Supplementary Table S1. All sequences are available from GenBank with accession numbers or gene locus tag numbers indicated in parentheses, including *P. jirovecii* (T551_00002), *P. macacae* (MN509821), *P. oryctolagi* (MN509824), *P. canis* (MN509823), *P. carinii* (T552_04149), *P. fulvescens* (MN509819), *P. murina* (PNEG_04309), *P. muelleri* (MN509817), *P. tanezumi* (MN509820), *P. wakefieldiae* (AF164562) and *P. exulans* (MN509818). UCSs in *P. murina, P. carinii, P. wakefieldiae* and *P. jirovecii* have been reported previously as detailed in the main text while all others were identified in this study.

The Msg-A2 subfamily represents the previously reported *msr* genes in *P. carinii* (24, 25) and differs from Msg-A1 as follows: 1) The abundant presence in rodent *Pneumocystis*, but absence in all other species examined thus far (Table 1); 2) The presence of a short intron near the 5’ end; 3) The presence of a unique, highly conserved exon 1; and 4) Independent expression of each individual gene (not under the control of UCS). Alignment of 73 full-length *msg*-A2 genes from rodent *Pneumocystis* revealed two groups of genes with a size of ∼2 kb and ∼3 kb, respectively. The group with a larger size contains all 9 Msg domains while the other group lacks three of them (M5, M6 and C1). In phylogenetic analysis, all *msg*-A2 genes from *P. carinii* form a strong clade (with 99% bootstrap support) while *msg*-A2 genes from *P. wakefieldiae* intersperse among *msg*-A2 genes from *P. murina* (Fig. S2). In *P. carinii*, 11 *msg*-A2 genes show higher sequence identities to *msg*-A1 genes compared to other *msg*-A2 genes (53-63% *vs* 35-44%), and are clustered together with *msg*-A1 genes from *P. carinii* in phylogenetic analysis (Fig. S3). Similarly, one of the 6 *msg*-A2 genes in *P. wakefieldiae* shows higher sequence identity to and is clustered with *msg-*A1 genes.

The Msg-A3 subfamily includes genes with substantial sequence identity to the Msg-A1 and Msg-A2 subfamilies but without the CRJE element of the *msg*-A1 genes or the highly conserved exon-1 of the *msg*-A2 genes (Fig. S4). This subfamily has a significant expansion in *P. jirovecii* with 33 copies but only 1 to 6 copies in other species (Table 1). With an overall highly variable 5’-end leader, members of this subfamily are expected to be expressed independently, similar to the Msg-A2 subfamily. Nevertheless, 5 of the 6 *msg*-A3 genes in *P. murina* contain a ∼600 bp leader sequence with significant identity and structural similarity to the UCS (termed UCS-like), including a relatively long intron (Fig. 4). These 5 genes are distributed in different chromosomes. Based on reverse-transcription PCR studies, each of these 5 genes was expressed independently (Text S1). Of note, the Msg-A3 subfamily encompasses both the Msg-II and Msg-III families reported by others (17), as illustrated in Fig. S5. Given the complex clustering patterns in phylogenetic analysis and unknown functions of these genes, it appears not meaningful to further divide the Msg-A3 subfamily.

**FIG 4.**
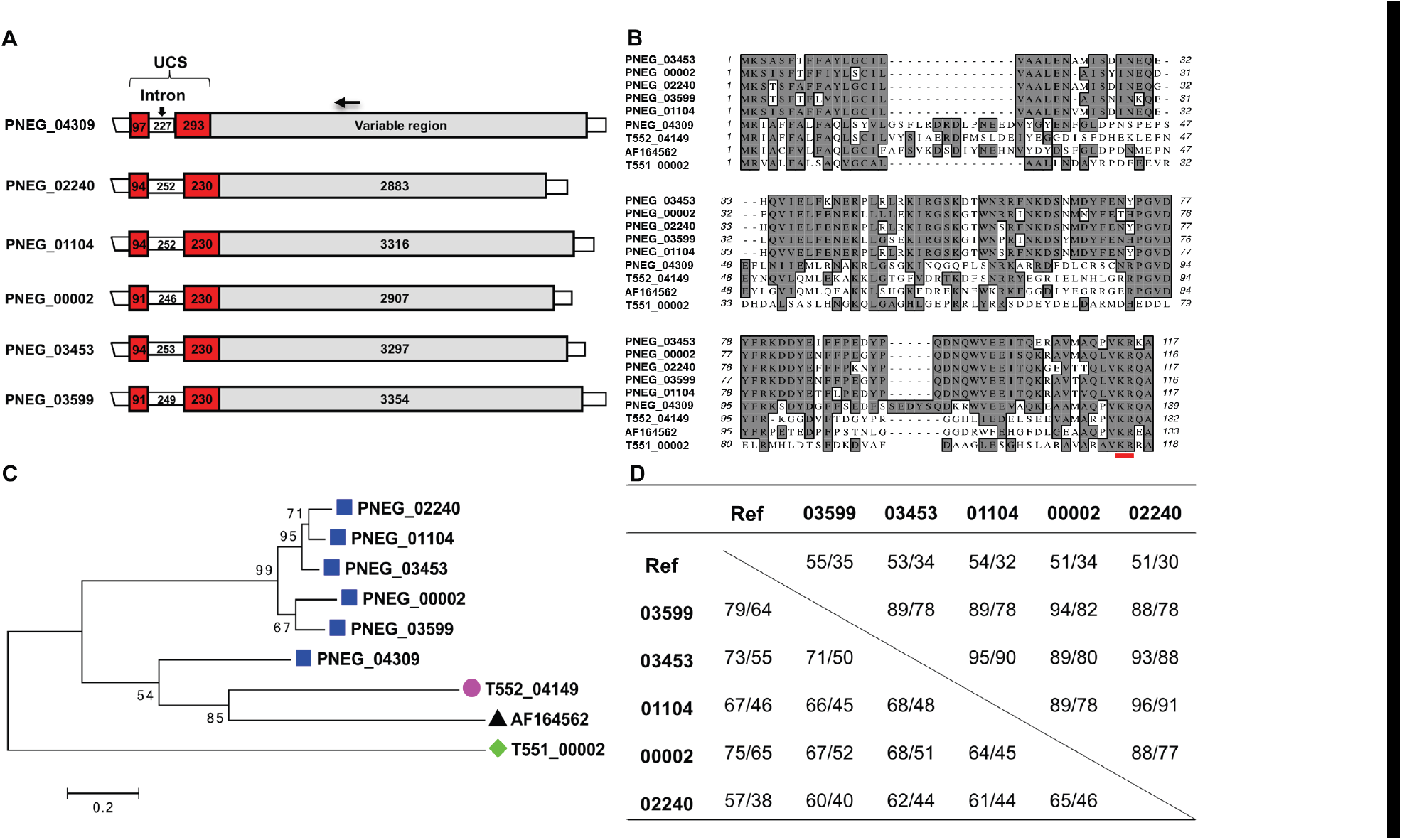
Five *msg*-A3 genes containing a UCS-like leader sequence in *P. murina*. **(A)** Schematic representations of *msg* genes, including 5 containing a UCS-like sequence (PNEG_02240, PNEG_01104, PNEG_00002, PNEG_03453 and PNEG_03599) and one classical *msg*-A1 gene (PNEG_04309). The numbers in the boxes represent the size (bp) of the regions indicated. The horizontal arrow on the top indicates the approximate location of the reverse primer MSG.r2b conserved among all *msg*-A1 and *msg*-A3 genes and used to determine the independent expression of the 5 *msg*-A3 genes (Table S2 and Text S1). (**B**) Alignment of the UCS and UCS-like protein sequences, including all those shown in panel A and the UCS in *P. carinii* (T552_04149), *P. wakefieldiae* (with GenBank accession code AF164562), and *P. jirovecii* (T551_00002). KR (red underlined) represents putative cleavage site by kexin-like endoprotease. (**C**) Phylogenetic relationship of UCS and UCS-like proteins based on sequences shown in panel B. Numbers on the branches indicate bootstrap support values. (**D**) Sequence identity among the *msg* genes shown in panel A (without including the first 4 characters of the gene IDs). Ref refers to the UCS gene PNEG_04309 and the *msg*-A1 gene PNEG_04309. Values in the table refer to the identity (%) of nucleotide and amino acid sequences in UCS (top right) and variable regions (lower left).

### Msg-B family

This represents the only Msg family completely absent in all rodent *Pneumocystis* sequenced to date, but with exceptionally high abundance in *P. jirovecii* (Table 1). With a highly variable 5’-end leader, members of this family are expected to be expressed independently. In phylogenetic analysis (Fig. S6), the family is separated into two major groups, which also differ in size (1.3 kb and 1.6 kb, respectively).

### Msg-C family

The prominent characteristics of this family are its significant presence in *P. murina* and unique chromosomal organization (Fig. S7). This family consists of a tandem array of 6 genes in chromosome 17 of *P. murina,* which represents the largest tandem array of genes of the same family identified so far in any *Pneumocystis* species. In contrast, there are no more than two *msg*-C genes in other *Pneumocystis* species. The two *msg-*C genes in *P. wakefiediae* share a similar size, intron-exon structure and domain compositions (N1, M2 and M3) with *msg*-C genes in *P. murina*. However, in all other species examined, the *msg*-C genes are smaller (0.8-1 kb) with different intron-exon structures, and/or lack the highly conserved exon-1 sequence of *P. murina*. In addition, they have different domain compositions and are only distantly related to the 6 genes in *P. murina* by phylogenetic analysis (Fig. S7). Furthermore, the chromosomal arrangement of the *msg*-C genes in *P. jirovecii* and *P. macacae* is different from that in rodent *Pneumocystis* (Fig. S7C). It is likely that the *msg*-C genes in *P. carinii, P. jirovecii* and *P. macacae* represent degenerate genes or pseudogenes, as supported by the low-level transcription of this gene in *P. carinii* (Table 2).

**Table 2.** Relative expression levels of the *msg* superfamily in *P. murina* and *P. carinii.*

| Genes | *P. murina* |  | *P. carinii* |  |
| --- | --- | --- | --- | --- |
|  | No. of genes | FPKM<sup>a</sup> | No. of genes | FPKM<sup>a</sup> |
| *msg*-A1 | 22 | 670 (15-2,011) | 65 | 141 (4-2,550) |
| *msg*-A2 | 14 | 252 (72-556) | 53 | 205 (0-1,202) |
| *msg*-A3 | 6 | 725 (77-2,015) | 3 | 112 (72-929) |
| *msg*-C | 6 | 3,895 (662-9,382) | 1 | 41 |
| *msg*-D | 1 | 1,385 | 1 | 2,154 |
| *msg*-E | 7 | 839 (100-4,270) | 5 | 1,742 (122-8,010) |
| Unclassified | 8 | 50 (0-90) | 13 | 25 (8-158) |
| UCS | 1 | 17,646 | 1 | 15,830 |
| Genome | 3,623 | 152 (0-17,646)<sup>b</sup> | 3,646 | 148 (0-15,830)<sup>b</sup> |

<sup>a</sup>Fragments per kb of exon per million fragments mapped based on RNA-Seq data as described in reference (1). Data are expressed as median (minimum-maximum) for multi-copy gene families.
<sup>b</sup>Median values for all protein-coding genes in the genome.

### Msg-D family

This family is related to the previously reported A12 antigen gene in *P. murina* (26). Like Msg-A3 subfamily and Msg-B family, this family is rarely present in rodent *Pneumocystis* but significantly expanded in *P. jirovecii* and perhaps in *P. macacae* as well (Table 1). However, most of the Msg-D members contain 6 conserved domains (Fig. S8) compared to 9 and 3 domains in Msg-A3 and Msg-B, respectively. In phylogenetic analysis, all single-copy Msg-D members in rodent *Pneumocystis* tightly cluster together into one clade, which is well separated from Msg-D members in all other species. Consistent with the phylogenetic analysis, Msg-D members in all rodent *Pneumocystis* lack both N1 and M2 domains, which are present in most of Msg-D members in other species.

### Msg-E family

This family is related to two previously reported p55 antigen genes (27-29). Unique among all Msg families, the Msg-E family is the smallest in member size, molecular size and number of conserved domains. It is relatively equally distributed across all *Pneumocystis* species examined (Table 1) and among the most highly expressed families in rodent *Pneumocystis* (Table 2). Members do not cluster by species in phylogenetic analysis (Fig. S9). In *P. murina* there are three members with a nearly identical sequence and molecular size (termed p57), which are located in separate chromosomes (1, 30). Each of these 3 genes is present as a tandem array with one *msg*-A2 gene and one *msg*-A1 gene located downstream (1). Homologs to these 3 genes are also present in three separate chromosomes of *P. wakefieldiae* though their downstream genes have not been identified presumably due to incomplete genome assembly. No other species sequenced to date have close homologs to these 3 genes. These findings further suggest that duplication of the p57 gene in *P. murina* and *P. wakefieldiae* occurred before separation of these two species, or there was introgression.

### Unclassified genes

In *P. murina, P. carinii* and *P. jirovecii*, there are 8 to 13 genes related to Msg that are unable to be reliably classified due to their shorter length (∼970 bp on average), presence of multiple introns, or lack of unique sequences (CRJE, KR site or conserved leader sequences) (Data Set 8). The shorter length in most of them is not due to incomplete sequencing as they are present within well covered contigs. Almost all of these genes in *P. murina* and *P. carinii* have a low expression level (Table 2), suggesting they are degenerate genes or pseudogenes.

### Highly variable expression levels among different *msg* families in *P. murina* and *P. carinii*

The relative expression level for each gene was estimated using RNA-Seq data from three heavily infected laboratory animals each for *P. murina* and *P. carinii* as previously described (1). RNA-Seq data indicate that all *msg* genes in *P. murina* and *P. carinii* are transcribed except for two unclassified *msg* genes in *P. murina* and 5 *msg*-A2 genes in *P. carinii* (Table 2). Strikingly, the UCS gene in both *P. murina* and *P. carinii* was the most highly expressed protein-coding gene of the whole genome (with their FPKM values being more than 100 times higher than the median expression level for the whole gene set); as expected, individual *msg*-A1 members were expressed at lower levels. This high expression level is consistent with SDS-PAGE analysis of *Pneumocystis* proteins, which demonstrates that Msg is the most abundant protein as estimated by Coomassie blue staining (1). In *P. murina*, the highest expression level of individual Msg genes was observed in the Msg-C family, followed by the Msg-D, Msg-E, Msg-A3 and Msg-A1 family or subfamily, all of which showed an expression level at least 3 times higher than the median of the whole gene set. The expression level of the Msg-A2 family was slightly higher than the median. In *P. carinii*, the highest expression level was observed in the single Msg-D gene, followed by the Msg-E family, both of which showed an expression level at least 11 times higher than the median. The expression level of the Msg-A family (including all 3 subfamilies) was similar to the median.

### Significant diversity of UCS in *Pneumocystis* from 10 mammalian host species

UCS has been previously reported for *P. murina, P. carinii, P. wakefieldiae* and *P. jirovecii* (19-22, 31). In the present study we identified UCS in *Pneumocystis* infecting rhesus macaques, dogs, rabbits, chestnut white-bellied rats, Müller’s giant Sunda rats, Asian house rats and Polynesian rats. Details about the nomenclature of these mammalian species and *Pneumocystis* species are listed in Table S1. As shown in Fig. 3, the *Pneumocystis* UCSs from all these animals show the sequence organization in known UCSs, including two exons that are interrupted by a variably sized intron. While exon 1 is identical in size (97 bp) among all UCSs, exon 2 is highly variable in size, with the shortest size present in *P. oryctolagi* (230 bp) and the longest in *P. murina* (314 bp).

The predicted UCS protein sequences vary in size from 110 to 138 amino acids, with 24-97% sequence identity (Fig. S11). Despite these variations, all UCSs contain a pair of basic amino acid residues in the carboxyl end, Lys-Arg, known as the KR site (20, 32, 33). Phylogenetic analysis showed a clear separation between the UCSs in *Pneumocystis* from rodents and those from other mammalian species (Fig. 3A). Consistent with the phylogenetic relationships, the UCSs in all rodent *Pneumocystis* had an extra 13-15 amino acid residues at the beginning of exon 2 and a unique hexapeptide of PGVDYF near the center of exon 2 compared to *Pneumocystis* from other mammalian species (Fig. S11).

Similar to exon 2, the intron is also highly variable in size, with the shortest present in *P. carinii* (150 bp) and the longest in *P. oryctolagi* (515 bp). In addition, different levels of inter- or intra-strain sequence variation were observed in UCSs from *P. carinii, P. wakefieldiae, P. macacae* and *P. oryctolagi* (Text S1 and Figs. S12-S13). The highest variation was observed in *P. oryctolagi* isolates, which displayed extensive inter- and intra-strain variations, including two SNPs in exon 2 and many SNPs, indels and tandem repeat variations in the intron (Fig. 5).

**FIG 5.**
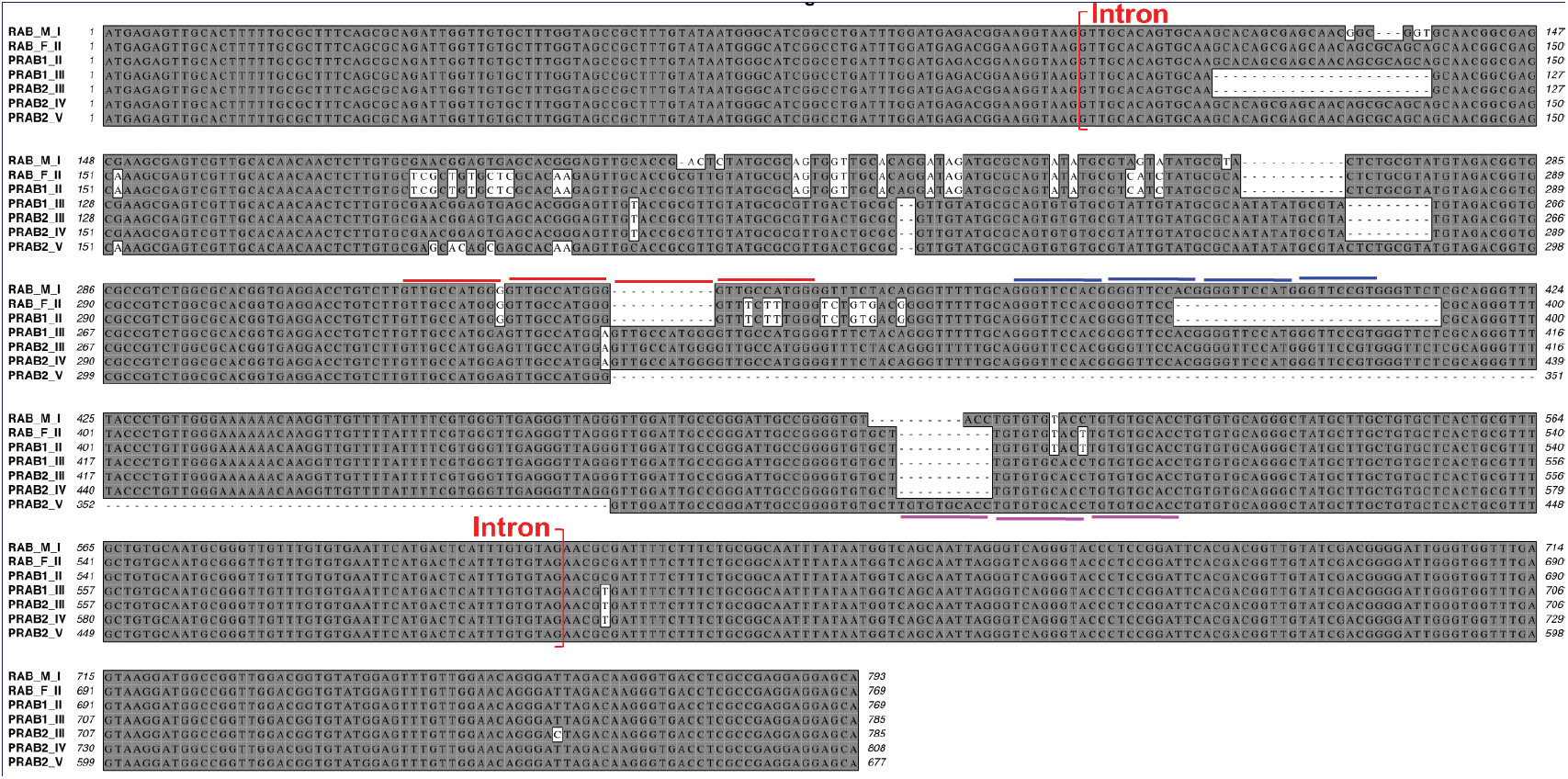
Sequence variation in the expression site (UCS) of the *msg*-A1 gene in 4 *P. oryctolagi* isolates. Five different sequence populations were identified, which are named as types I to V indicated at the end of the sample codes, including RAB_M from Michigan, USA; RAB_F from Tours, France; and PRAB1 and PRAB2 from Lille, France. The 3 types of sequences (III, IV and V) in sample PRAB2 were obtained from sequencing of 2, 6 and 3 plasmid clones, respectively, while the 2 types of sequences (II and III) in sample PRAB1 were obtained from 3 and 5 plasmid clones, respectively. The other 2 samples showed no variation based on Illumina sequencing of genomic DNA; their PCR products showed homogeneous sequences and were not subjected to subcloning. Three types of tandem repeats are indicated by colored lines. Numbers at either side of the alignment refer to the nucleotide positions relative to the predicted UCS translational start site. The intron is indicated in red brackets. Sequences are available from GenBank with accession numbers MN509824-MN509828.

### Substantial variation of the *msg*-A1 gene repertoires in *P. carinii* from laboratory and wild rats

To compare the *msg* diversity between *Pneumocystis* from laboratory-bred animals and that from wild animals, we analyzed the restriction fragment length polymorphism (RFLP) patterns of the *msg-*A1 repertoires in *P. carinii* from 8 wild Norway rats collected in Ontario, Canada in comparison with *P. carinii* from 8 laboratory Norway rats collected in three different animal facilities in USA. *P. carinii* from all laboratory rats showed an almost identical RFLP pattern while substantial variations in the RFLP patterns were observed within *P. carinii* isolates from wild rats and between *P. carinii* from wild and laboratory rats (Fig. 6)

**FIG 6.**
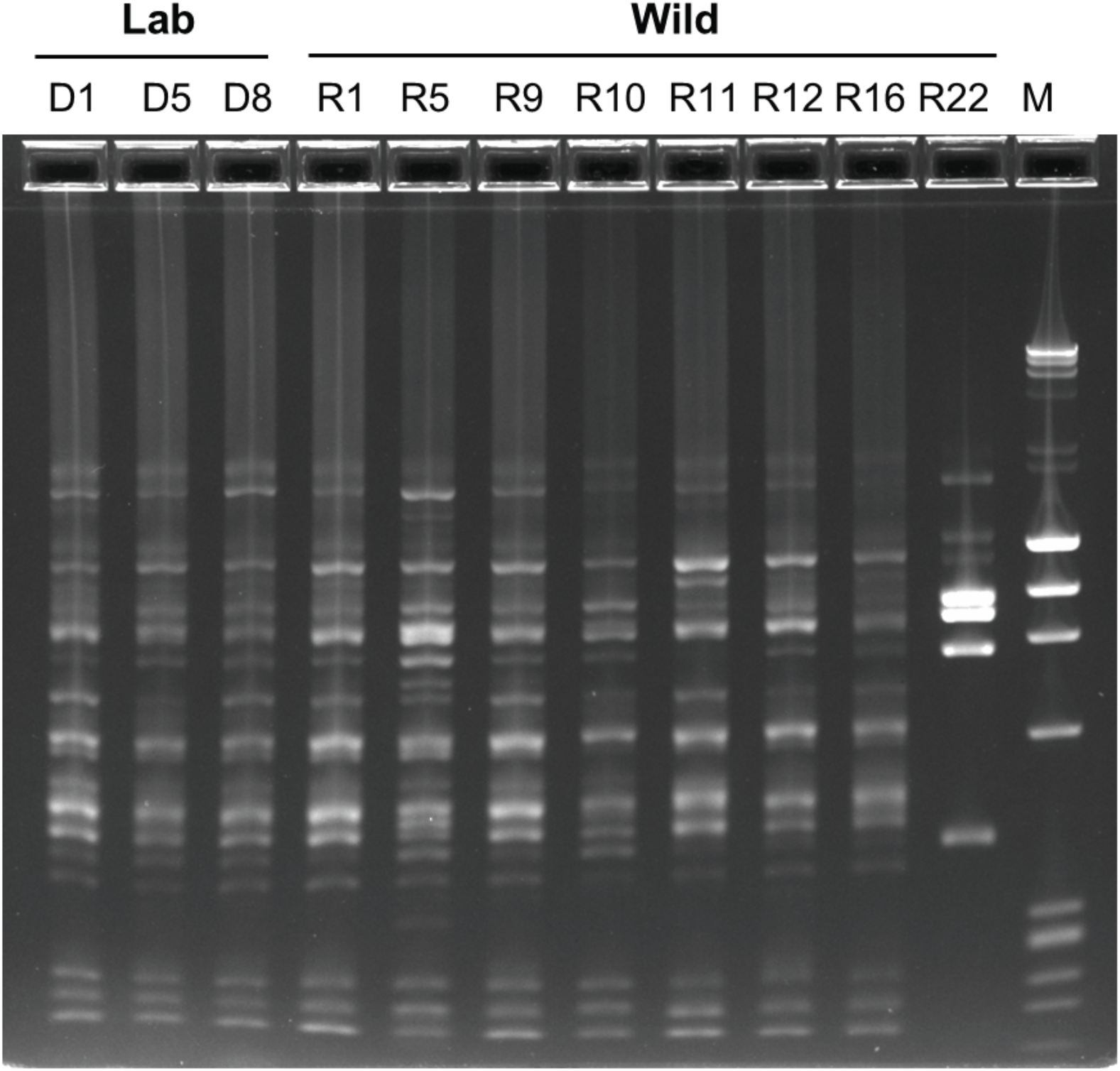
Comparative restriction-fragment length polymorphism (RFLP) analysis of *msg-*A1 of *P. carinii* infecting laboratory and wild Norway rats. *msg-*A1 repertoire was amplified by PCR with genomic DNA prepared from *P. carinii*-infected lung samples from 8 immunosuppressed laboratory Norway rats (with three representatives show in lanes indicated as Lab) and 8 wild Norway rats (in lanes indicated as Wild). While R22 is most clearly representative of a different RFLP pattern, more subtle differences are also apparent in some of the other wild rats (e.g. R5 and R11) compared to laboratory rats. The PCR products were digested with restriction enzyme *Dra*I, and separated in 2% agarose gels containing SYBR Safe. Lane M, DNA size marker containing λDNA digested with *Hind III* and ØX174 DNA digested with *Hae III.*

To further confirm these variations, we determined the full-length *msg*-A1 sequences in *P. carinii* from one wild rat by Sanger sequencing of cloned PCR products. Sequence analysis of 28 random clones identified 13 unique *msg* sequences, with an identity of 78-96% at the nucleotide level and of 63-95% at the amino acid level. All but 1 of these 13 genes were clearly different from the 65 *msg*-A1 genes of *P. carinii* from laboratory. In phylogenetic analysis, *msg-* A1 genes from wild and laboratory rats were interspersed (Fig. 2). These findings suggest a closely related but clearly distinct repertoire of *msg*-A1 genes in *P. carinii* from wild and laboratory rats.

## DISCUSSION

Over the past several decades, Msg has been the most extensively studied molecule in *Pneumocystis*, primarily due to its abundance, its role in pathogen-host interactions and its potential as a target for diagnosis of *Pneumocystis* infection. In this report, we present an in-depth analysis of the *msg* domain structure and the characteristics of each individual *msg* family or subfamily, including new *msg* genes identified from *P. oryctolagi, P. macacae, P. canis, P. wakefieldiae* and *P. carinii* infecting wild rats. The results from our analysis demonstrate a much greater complexity to this superfamily than was previously appreciated, expand the understanding of the primary structure, organization, phylogeny and expression patterns of the Msg superfamily, and provide a comprehensive basis for further investigation of the role of the Msg superfamily in *Pneumocystis* biology.

The Msg superfamily, particularly in *P. jirovecii* (179 members), represents the largest surface protein family identified to date in the fungal kingdom (34), which is surprising for an organism whose genome size is the smallest in the fungal kingdom sequenced to date, after the intracellular Microsporidia (35). *msg* genes are unique to *Pneumocystis* and account for 3-6% of the total genome, suggesting a critical role in the organism’s survival (1). The vast majority of *msg* genes are clustered in subtelomeric regions, which are presumably advantageous to foster DNA recombination and antigenic variation as has been found for surface protein genes in other pathogens (36). Their positioning is consistent with the notion that subtelomeric regions are favorable locations for fungal pathogens to acquire novel genes and foster evolution (37, 38).

By domain structure, phylogenetic analysis and expression control mechanisms, we have been able to classify the Msg superfamily into discrete families and subfamilies. Our classification based on exhaustive cataloging of *msg* genes from multiple *Pneumocystis* species is more comprehensive than the one described in a recent report, which was based on a limited number of *msg* genes from *P. jirovecii* alone (17). Thus, despite the consistency of four families/subfamilies between these two systems (Msg-A1, -B, -D and –E *vs* Msg-I, -IV, -V and – VI), two families (*msg-*A2 and *msg*-C), which are almost exclusively present in rodent *Pneumocystis,* are absent in the recent report (17). We also did not subdivide the *msg*-A3 subfamily due to the complex clustering patterns in phylogenetic analysis (Fig S4) and unknown functions of these genes. This classification will likely be refined when our understanding of the function of Msg is improved and this superfamily is better characterized for other *Pneumocystis* species.

Based on our analysis, there is substantial conservation among most Msg families or subfamilies across different *Pneumocystis* species, but there are also species-specific expansions or contractions. Among the three *Pneumocystis* species with the most complete dataset, *P. murina* has the fewest number of genes in the Msg superfamily while *P. jirovecii* has the most. These differences may be related to the larger body and therefore lung size and the need for a higher degree of antigenic variation to avoid immunologic memory in individual human hosts, who have a longer life-span than rodent hosts. The larger size of the Msg superfamily in *P. jirovecii* is attributable in part to the expansion of the classic Msg-A1 subfamily, as well as other families (including Msg-A3, Msg-B and Msg-D), which have no or limited representation in rodent *Pneumocystis*. Of note, *P. murina* possesses a set of 6 *msg* genes (Msg-C family) that are clustered as a tandem array in one chromosome and are the most highly expressed *msg* genes (Table 2).

The functions of Msg remain unknown or poorly understood. To date, the best studied genes of the Msg superfamily are those classical Msg genes in the Msg-A1 family, whose expression is regulated by the single-copy UCS expression site, which allows antigenic variation through DNA recombination (14, 17). Such variation can potentially serve as a mechanism to facilitate evasion of host immune responses, enabling the organism to persist longer in the host and transmit to a new host. This mechanism is presumably operational only in immunocompetent hosts. The expression of all msg-A1 variants in a given lung is presumably a consequence of immunodeficiency. For all three *Pneumocystis* species with nearly fully sequenced genomes, the *msg*-A1 genes account for approximately 50% of all *msg* genes, supporting their potential to efficiently generate a large number of variants allowing immune evasion. In support of this hypothesis, our RNA-Seq analysis of *P. murina* and *P. carinii* revealed an exceptionally high-level expression of UCS and a variable level expression of all individual *msg*-A1 genes (Table 2).

UCS is known to have a highly variable number of tandem repeats in the intron in *P. jirovecii* (20, 39, 40). In this study, we demonstrated for the first time the presence of inter- and intra-strain variations in tandem repeats in the intron of UCS in *P. carinii* and *P. oryctolagi.* UCS appears to have a higher degree of variation in tandem repeats as well as SNPs compared to *P. jirovecii* UCS. The intron in UCS (150-515 bp) is among the longest introns in *Pneumocystis* species studied to date. The retention of such a long intron with high variability in these species in an otherwise highly reduced genome suggest a critical role in organism survival, e.g. transcriptional regulation by a recursive splicing mechanism (41, 42).

Of note, while UCS is present as a single copy gene per genome in all *Pneumocystis* species, there are 5 *msg*-A3 genes in *P. murina*, each of which contains an UCS-like leader sequence (Fig 3) and is expressed at a relatively high level independent of the classic UCS (Table 2). These may have arisen from gene duplication in *P. murina*; alternatively, it is possible that a common ancestor of *Pneumocystis* had multiple UCSs, which have been gradually lost as a result of evolving an efficient recombination system involving only a single UCS (for the *msg*-A1 family),

Previous studies have demonstrated a conservation of the *msg*-A1 repertoires in *Pneumocystis* in colony-bred laboratory rats and mice in contrast to the highly variable *msg* repertoires in *P. jirovecii* (14), suggesting a homogeneous population of rodent *Pneumocystis* due to closed breeding conditions. In support of this, we observed substantial variations in the RFLP patterns within *P. carinii* isolates from wild rats and between *P. carinii* from wild and laboratory rats, supporting the former possibility. These variations may reflect the difference in immune system development and modulation in wild animals as they are continuously exposed to high levels of immune challenges in an open environment and experience high levels of infection with a wide range of pathogens (43-45). We hypothesize that this diversity is driven in part by a need for antigenic variation in response to T cell rather than B cell mediated immune responses, and potential adaptation to the diverse HLA repertoire that would be present in a natural community of host species (46) versus the limited diversity present in inbred communities.

Domains M1 to M5 of *msg*-A1 encoded proteins likely arose by gene duplication given their conserved pattern of cysteine residues, and in fact only a single M domain is categorized in PFAM. However, more detailed analysis clearly allows the identification of 5 related but unique domains. It is noteworthy that by phylogenetic analysis, individual domains are more closely related to each other across species than to other M domains in the same species, which suggests that there is a critical function for each domain and its evolution is restricted by negative selection. Further, given that *msg*-A1 encoded proteins with these domains have been identified in all *Pneumocystis* species studied to date, this gene organization appears to have developed in an ancestor common to all *Pneumocystis* species, and may have been a critical factor that allowed *Pneumocystis* to successfully infect mammalian hosts.

Unlike *P. jirovecii,* and perhaps *P. macacae, P. canis* and *P. oryctolagi*, rodent *Pneumocystis* (*P. murina, P. carinii* and *P. wakefieldiae*) have a large number of *msg*-A2 genes, which are only slightly less frequent than *msg*-A1 genes. Previous studies of *P carinii* have shown that *msg*-A2 genes are expressed independent of the UCS (24, 25). Nevertheless, the possibility of homologous recombination between *msg*-A2 and *msg*-A1 genes cannot be ruled out due to their high sequence identities, as previously suggested (13, 47). Eleven *msg*-A2 genes in *P. carinii* show higher identities to *msg*-A1 genes than to other *msg*-A2 genes in this organism. It is likely that these 11 *msg*-A2 genes (the second exons) have arisen as a result of reciprocal recombination with *msg*-A1 genes (through a mechanism unrelated to UCS or CRJE). While it appears that *msg*-A2 expression is not regulated by UCS, nothing is known yet about what mechanisms control the *msg*-A2 expression, or whether the *msg*-A2 family contributes to antigenic variation in response to immune pressure, environmental changes, or life cycle phases. The presence of a long monoguanosine repeat in some *msg*-A2 genes has raised the possibility that variation in the length of this repeat may cause frameshifts, thus altering the amino sequence downstream of the repeat (13, 47). However, based on the high-throughput genome sequencing data with at least 150 X coverage (1), sequence reads for the monoguanosine repeat region in all involved *msg*-A2 genes appeared highly uniform though a small number of reads (< 5%) showed different numbers of repeats. We could not determine if this was caused by sequence errors or *in vivo* changes. The presence of such a small number of variable reads does not seem to support an involvement of this repeat in altering the antigenicity or other functions of the *msg*-A2 genes. Of note, a polyguanosine repeat encodes a polyglycine peptide, which has been shown in other organisms to play various critical roles, such as protein-to-protein interactions, cell wall plasticity, and modulation of developmental stages (48-50). Whether the polyglycine peptide in Msg-A2 proteins has these functions awaits future investigation.

Despite their potential importance in *Pneumocystis*’ survival, the functions of the vast majority of members of the *msg* superfamily remain poorly understood or uncharacterized. Even for the most extensively studied *msg*-A1 genes, while it has been generally believed that their primary function is to confer antigenic variation and immune evasion, there are only limited experimental data supporting this potential function (46). There are also multiple studies showing an involvement of Msg proteins in mediating adherence of *Pneumocystis* organisms to host alveolar epithelial cells and macrophages (51-53) though it is unknown if the Msg proteins involved in these studies represent Msg-A1 or other Msg proteins, especially Msg-A2 and Msg-A3 proteins, which are highly similar to Msg-A1 proteins in sequence and length. The functions of all non-*msg*-A1 genes remains unknown. They may provide other advantages for *Pneumocystis* to survive in the host, such as mediating developmental states, optimizing mobility and adhesion ability, and adapting to specific host niches or environmental conditions. In support of this hypothesis, one such gene of the *msg*-E family in *P. murina*, termed p57, has been shown to be a stage-specific antigen that is expressed exclusively on intracystic bodies and young trophic forms, suggesting a role in the *Pneumocystis* life cycle development (30).

In conclusion, despite a highly reduced genome, *Pneumocystis* is equipped with a large complex superfamily of *msg* genes. These genes exhibit conservation among *msg* families and subfamilies across different *Pneumocystis* species, as well as species-specific expansions or contractions. The versatility of these genes may mirror their association with a wide variety of functions, rather than just conferring antigenic variation to allow immune evasion as previously believed. Our results provide a rich source of information that lays the foundation for the continued experimental exploration of the function of the Msg superfamily in *Pneumocystis* biology.

## METHODS

### Sources of *Pneumocystis msg* sequences

The primary source of *msg* sequences for *P. murina, P. carinii* and *P. jirovecii* was from our previous studies (1, 15, 18), which are available from the NCBI Umbrella project PRJNA223519 at http://www.ncbi.nlm.nih.gov/bioproject/?term=PRJNA223519. In this study we obtained additional *Pneumocystis msg* and UCS sequences from various animals as listed in Table S1, which includes new, tentative names for *Pneumocystis* organisms not reported previously; names are based on the rules of the MycoBank Database (http://www.mycobank.org) for new fungal names. All new sequences determined in this study are available from the NCBI BioProject database with accession no. PRJNA560924. The methods to obtain these sequences are described below.

### *Pneumocystis* sample sources and DNA extraction

Agarose gel blocks containing *P. wakefiediae* and *P. carinii* were obtained from 4 Norway rats immunosuppressed once per week by 4 mg/kg of DepoMedrol (Pharmacia and Upjohn Co. a division of Pfizer, Inc.) at the animal facility of the University of Cincinnati, Ohio, USA (54). Genomic DNA in gel blocks was extracted using the Zymoclean Gel DNA Recovery Kit (Zymo Research).

*P. carinii*-infected lung tissues were obtained from 8 immunosuppressed Sprague Dawley rats collected between 1986 and 2013 from the animal facilities at NIH, Bethesda, Maryland (14, 55); Indiana University, Indianapolis, Indiana (56); and Louisiana State University Health Science Center, New Orleans, Louisiana. Genomic DNA was isolated by use of either a QIAamp DNA Mini kit (Qiagen) or a traditional method utilizing proteinase K digestion, phenol-chloroform extraction and ethanol precipitation (14). In addition, *P. carinii*-infected lung tissues were obtained from 8 wild Norway rats (*R. norvegicus*) from pig farms in Ontario, Canada in 2015 as previously described (57). Genomic DNA was extracted using the MasterPure Yeast DNA Purification Kit (Epicentre). All these wild rats appeared to be healthy upon capture and were confirmed to be infected by *P. carinii* alone based on sequence analysis of two *Pneumocystis* mitochondrial genes (large subunit of rRNA and ATPase subunit 6. Unpublished data).

DNA samples for *Pneumocystis* infecting other wild rat species in Southeast Asia, including chestnut white-bellied rats, Müller’s giant Sunda rats, Asian house rats and Polynesian rats, were obtained from previous studies (58). All animals appeared to be healthy upon capture.

*P. macacae*-infected lungs were obtained from two SIV-infected rhesus macaques at the NIH Animal Center, Bethesda, Maryland, USA (59, 60). Genomic DNA was extracted following a *Pneumocystis* DNA enrichment protocol as described previously (1). An additional two *P. macacae* samples were obtained as formalin fixed paraffin embedded (FFPE) tissue sections prepared from SIV-infected rhesus macaques at the Tulane National Primate Research Center, Covington, Louisiana (61) and the UC Davis California National Primate Research Center, Davis, California, USA, respectively. Genomic DNA was extracted using the AllPrep DNA/RNA FFPE Kit (Qiagen).

*P. canis* DNA samples were obtained from one Cavalier King Charles Spaniel dog at the University of Helsinki, Finland (62, 63) and one Whippet dog at the University of Veterinary Medicine, Vienna, Austria (64), respectively.

Four *P. oryctolagi* DNA samples were obtained from previous studies of one rabbit with severe combined immunodeficiency at the University of Michigan, Ann Arbor, Michigan, USA (65), and three immunosuppressed rabbits at the Institut Pasteur de Lille (66) and the Institut National de la Recherche Agronomique de Tours Pathologie Aviare et Parasitologie, Tours (67), France.

Animal experimentation guidelines of the National Institutes of Health were followed in the conduct of these studies.

### Illumina sequencing

DNA extracts for 4 *P. wakefieldiae*, 4 *P. macacae*, 2 *P. canis* and 4 *P. oryctolagi* samples were subjected to whole genome sequencing commercially in an Illumina HiSeq platform using a 150-bp paired-end library and/or a 250-bp paired-end library. Genome assembly was performed essentially as previously described (1, 68); detailed analyses of these genomes will be published separately.

### *msg* sequences of *P. wakefiediae*

To amplify the repertoire of the classical *msg*-A1 genes in full length, the forward primer (WSG.f3) was designed from the 3’-end of the previously reported UCS (within CRJE) of *P. wakefiediae* (GenBank accession no. AF164562) (31). The reverse primer (WSG.r5) was designed from the highly conserved 3’-end coding region near the stop codon based on an alignment of more than 3,000 Illumina HiSeq reads. Primer sequences are listed in Table S2. Both primers were specific for *P. wakefiediae,* with no cross amplification of *P. carinii*. PCR was performed using *P. wakefiediae* genomic DNA and the AccuPrime Pfx SuperMix kit (Thermo Fisher Scientific) with the following cycling conditions: 95°C for 5 min and then 35 cycles at 95°C for 15 sec, 55°C for 30 sec, and 68°C for 3 min, with a final extension at 68°C for 5 min. The PCR product was subcloned into pCR2.1 TOPO vector by use of the TOPO TA Cloning kit (Invitrogen, Carlsbad, CA). Two clones containing the full-length *msg*-A1 gene were sequenced commercially by Sanger sequencing.

To sequence the *P. wakefiediae* homologue of the 6-gene cluster of the *msg*-C family in *P. murina*, we first used the Illumina reads of *P. wakefiediae* (mixed with *P. carinii* reads) to assemble the *P. wakefiediae* homologues of PNEG_03432 and PNEG_03438, which are flanking the 6-gene cluster in *P. murina*. Subsequently, we designed a primer set (3432.f1 and 3438.r1. Table S2) specific for these two genes in *P. wakefiediae*. With these two primers, we amplified an 8-kb fragment from *P. wakefiediae* DNA and sequenced its full-length by Sanger sequencing with primer-walking. From a draft *P. wakefiediae* genome assembly we identified members of the *msg*-A2, *msg*-A3, *msg*-D and *msg*-E families or subfamilies based on homology to known genes in *P. murina, P. carinii* and *P. jirovecii* (1). Full-length *msg*-A1 genes sequences were unable to be assembled from the short HiSeq reads (16).

### *msg*-A1 sequences of *P. carinii* from wild rats

To determine whether the *msg-*A1 repertoires are identical in *P. carinii* from wild and laboratory Norway rats, we performed RFLP analysis of *P. carinii* isolates from 8 wild rats in comparison with 8 laboratory rats. The *msg*-A1 repertoires were amplified by PCR using primers RSG.f10 and RSG.r8 (Table S2), which are located in the highly conserved regions at the beginning and end of the *msg* coding regions, respectively, among 57 known full-length *msg-*A1 genes in *P. carinii* (1). Amplification was performed using the LongAmp Master Mix (New England Biolabs) with the follows parameters: 94°C for 2 min and then 35 cycles at 94°C for 15 sec, 55°C for 30 sec, and 68°C for 3 min, with a final extension at 68°C for 5 min. PCR products were purified and subjected to restriction digestion with DraI (New England Biolabs) at 37°C for 2 h. The resulting digests were purified and separated in 2% E-gel containing ethidium bromide (Invitrogen, Carlsbad, CA).

The *msg*-A1 repertoire from one wild rat (no. R5), which showed a distinct RFLP pattern compared to laboratory rats, was chosen for sequencing after PCR amplification using the primer pair RSG.f10-RSG.r8 and the LiSpark Max SuFi PCR Master Mix kit (LifeSct LLC, Rockville, MD). The PCR product was subcloned into pCR-XL-2 TOPO vector by use of the TOPO XL-2 Complete PCR Cloning kit (Invitrogen, Carlsbad, CA). A total of 28 clones containing the full-length *msg*-A1 gene were sequenced commercially by Sanger sequencing.

### *msg* sequences of *P. macacae, P. canis* and *P. oryctolagi*

Illumina HiSeq reads from one *P. macacae* sample were aligned to all known full-length *msg*-A1 genes of *P. jirovecii* (1), resulting in at least 1,000 reads for the very 5’- and 3’-end of the *msg*-A1 coding regions, respectively. Two primers (KSG.f3 and KSG.r2. Table S2) were designed from highly conserved regions based on alignment of these reads. The full-length *msg-*A1 repertoire in *P. macacae* was amplified using these two primers and the LiSpark Max SuFi PCR Master Mix kit, followed by subcloning into the pCR-XL-2 TOPO vector as described above. Two clones containing the full-length *msg*-A1 gene were sequenced commercially by Sanger sequencing.

For other *msg* families and subfamilies, we identified a small number of representative genes from a partial genome assembly of *P. macacae* based on homology to known genes in *P. murina, P. carinii* and *P. jirovecii* (1).

For *P. canis* and *P. oryctolagi,* a small number of genes representing each *msg* family were identified from a partial genome assembly of *P. canis* and *P. oryctolagi*, respectively.

### UCSs of *msg*-A1 genes in *Pneumocystis* from various mammalian host species

The UCS and its 5’-UTR sequences in *P. macacae, P. canis* and *P. oryctolagi* were first obtained by assembling Illumina HiSeq reads from whole genome sequencing as described above, followed by confirmation by PCR amplification and Sanger sequencing of genomic DNA. Based on sequence alignment of these UCSs and know UCSs of *P. murina, P. carinii, P. wakefieldiae* and *P. jirovecii*, we designed one forward primer (5UTR) from the conserved region in 5’-UTR and one reverse primer (CRJE.r3) from the conserved region in CRJE (Table S2). This primer set was used to amplify the UCS along with its 5’-UTR in *Pneumocystis* from other mammal species including dogs, rabbits, chestnut white-bellied rats, Müller’s giant Sunda rats, Asian house rats and Polynesian rats (Fig. 4 and Table S1). To study the variability of UCS and its downstream *msg-*A1 coding region in different *P. oryctolagi* isolates, PCR was performed using a pair of primers OSG.f3 and OSG.r9, which are located at the very 5’ end of UCS and one highly conserved region near the 5’ end of the *msg*-A1 coding region (Table S2). All PCR products were sequenced directly and/or after subcloning into the pCR2.1 TOPO vector as described above.

### Phylogenetic analysis

To analyze phylogenetic relationships, deduced protein sequences were aligned using MUSCLE (69) and phylogenetic trees were constructed based on maximum likelihood (ML) using RAxML (v8.2.5) (70) with 100 bootstraps as support values. The best amino acid model was estimated using PROTGAMMAAUTO option.

## Supporting information

Supplemental Data Set 1

Supplemental Data Set 2

Supplemental Data Set 3

Supplemental Data Set 4

Supplemental Data Set 5

Supplemental Data Set 6

Supplemental Data Set 7

Supplemental Data Set 8

## Acknowledgements

This study has been funded in whole or in part with federal funds from the Intramural Research Program of the U.S. National Institutes of Health Clinical Center; the National Institute of Allergy and Infectious Diseases; the National Cancer Institute, National Institutes of Health, under Contract No. HHSN261200800001E; the National Human Genome Research Institute (grant U54HG003067 to the Broad Institute); the National Institute of Diabetes & Digestive & Kidney Diseases (grant R01DK109883 to Tulane National Primate Research Center); and the Office of Research Infrastructure Programs/OD (award P51OD011107 to CNPRC). The content of this publication does not necessarily reflect the views or policies of the Department of Health and Human Services, nor does mention of trade names, commercial products, or organizations imply endorsement by the U.S. Government. We thank Mr. Rene Costello for providing animal care, Dr. Nicolas Cere (Institut National de la Recherche Agronomique de Tours Pathologie Aviare et Parasitologie, Tours, France) for kindly providing *P. oryctolagi* samples, and B. Scandrett, C. Roehrig, K. Konecsni and participating farmers for coordinating rat sample collection in Ontario. We also thank Dr. Phillippe Hauser for providing information about *msg* sequences in their studies (17). The authors declare no conflicts of interests.

