## Supplemental Text S1, Tables S1 and S2, and Figures S1 to S13 for "Diversity and complexity of the large surface protein family in the compacted genomes of various *Pneumocystis* species"

**Ma *et al*.**

SUPPLEMENTAL TEXT S1

Distribution of *msg* superfamily genes in genome assemblies…………………………..…2
Characterization of conserved domains in Msg proteins…………………………………....2
Amino acid usage in Msg proteins…………………………………………………………..3
mRNA expression of *msg*-A3 genes containing UCS-like sequences………………………..4
Sequence variation of UCS in different *Pneumocystis* species………………………………..5

SUPPLEMENTAL TABLES

Table S1 Nomenclature of *Pneumocystis*………………………………………………..6
Table S2 PCR primers……………………………………………………………………..7

SUPPLEMENTAL FIGURES

Fig. S1 Phylogeny of Msg domains ……………………………………………..………8
Fig. S2 Phylogeny and domains of Msg-A2 family……………………………..………9
Fig. S3 Phylogeny of Msg-A1 and Msg-A2 families…………………………...……10
Fig. S4 Phylogeny and domains of Msg-A3 family……………………………......….11
Fig. S5 Comparison of Msg-A1 and Msg-A3 subfamilies from different studies……..……12
Fig. S6 Phylogeny and domains of Msg-B family……………………………..……..13
Fig. S7 Phylogeny and domains of Msg-C family……………………………..……..14
Fig. S8 Phylogeny and domains of Msg-D family……………………………..……..16
Fig. S9 Phylogeny and domains of Msg-E family……………………………..……..17
Fig. S10 Amino acid usage in Msg proteins……..…………………………..…………18
Fig. S11 UCS proteins from different *Pneumocystis* species…………………………..…..19
Fig. S12 UCS variation in *P. wakefieldiae*……………………………..……..……..….20
Fig. S13 UCS variation in *P. carinii*………………………………..…………..………..21

SUPPLEMENTAL REFERENCES

SUPPLEMENTAL REFERENCES……………………………..…………………………….22

SUPPLEMENTAL DATASETS (for MSG sequences) in 8 separate Excel files

**Supplemental Text S1**

**Distribution of *msg* superfamily genes in genome assemblies of *P. murina*, *P. carinii* and *P. jirovecii***

Of the 64 *msg* genes identified in *P. murina* (1), 61 were distributed in 17 large scaffolds (corresponding to chromosomes) and the remaining 3 were each in a small contig (~3 kb). Of the 61 genes in 17 large scaffolds, 52 were located in subtelomeric regions and the remaining 9 were internal. There were 12 clusters each containing two *msg* genes, 7 clusters each containing three genes, and one cluster containing 6 *msg* genes. There were three sets of duplicated clusters (with 95-99% identity), each containing two or three *msg* genes.

Of the 141 *msg* genes identified in *P. carinii* (1), 54 were distributed in 16 of the 17 large scaffolds (corresponding to chromosomes) and the remaining 87 were distributed among 81 small contigs. Of the 54 genes in 16 large scaffolds, 51 were located in subtelomeric regions and the remaining 3 were internal. There were a total of 11 clusters each containing two to seven *msg* genes, with *kex* genes present in 6 clusters. Two of these clusters, each containing four *msg* genes and two *kex* genes, appeared to be the result of duplication, with 99% identity. *Kex* genes, like *msg*, are members of a multicopy gene family predicted to encode surface proteins, that appear to be the result of duplication of a serine endoprotease, which localizes to the Golgi apparatus in other fungi. Multiple *kex* genes are found only in *P. carinii*; *P. murina* and *P. jirovecii* have only a single *kex* gene.

Of the 179 *msg* genes identified in *P. jirovecii* (1), 98 were distributed in 20 large scaffolds and the remaining 81 were distributed among 51 small contigs. All 98 genes in the large scaffolds were located in the ends of scaffolds, presumably representing subtelomeric regions. There were a total of 24 clusters, each containing two to nine *msg* genes.

The location of *msg* genes in other *Pneumocystis* species was not determined in this study due to incomplete genome assemblies.

**Characterization of conserved domains in Msg proteins**

Currently, there are only two domains of Msg described in the PFAM database (https://pfam.xfam.org): MSG (PF02349) and Msg2_C (PF12373), which are derived from a small number of *msg* sequences identified in *P. murina* (2) and *P. jirovecii* (3, 4). Classic Msg proteins typically have 5 separate MSG domains (named M1 to M5) and one Msg2_C domain (named C1). In order to refine the Msg domain categorization, and to determine if there were unique characteristics for each of these domains, we focused on Msg proteins from *P. murina*,

*P. carinii* and *P. jirovecii* (5) that were greater than 900 amino acid residues. Based on the coordinates identified by Pfam domain analysis, we extracted the sequence regions corresponding to the 5 MSG domains (M1 to M5). A tree built from the extracted domain regions demonstrates that these 5 domains are well separated phylogenetically (Fig. S1).

A detailed examination of the alignment of Msg proteins greater than 900 amino acids identified three additional conserved regions (named N1, M6 and C2) in addition to the M1 to M5 and Msg2_C (C1) domains (Fig. 1). To be able to identify other proteins with these domains, we built separate HMM models for each of these domains (available from the Zenodo database at https://zenodo.org/record/3515473#.Xa4nhCV7nUY). We then used all these 9 HMM models (N1, M1-M6, and C1, C2) to scan all available *Pneumocystis* proteins greater than 300 amino acids, to identify additional Msg related genes. The resulting top significant hit was used to rebuild each model. After three rounds of rebuilding, we generated the revised models that led to the identification of members of the Msg superfamily as shown in our previous report (1). Each of the revised M1 to M5 domains has 7 or 8 cysteine residues, which are regularly spaced and highly conserved among these five domains. Each of the other four domains (N1, M6, C1 and C2) has 2 to 5 cysteine residues that are again highly conserved. Our revised HMM models had higher sensitivity in detecting conserved regions in Msg members compared to the previous Msg domain models in Pfam database; some of the genes identified as Msg superfamily members using the newer HMM models were not identified by the Pfam analysis.

Each Msg family contains a different number of Msg domains, varying from 1 to 9 domains (Fig. 2 and Figs. S2 to S9). Domains N1 and M1-M5 are widely distributed among most families while domains M6, C1 and C2 are primarily present in the Msg-A family.

**Amino acid usage in Msg proteins**

Amino acid compositions were calculated using full-length Msg proteins in three Pneumocystis species with nearly fully sequenced genomes (*P. murina*, *P. carinii* and *P. jirovecii*). Overall, a bias towards Lys, Leu, Glu, Ser and Thr was observed in all 5 Msg families (Fig S10). In addition, there is a bias of amino acid usage in some particular regions of the Msg protein. Strikingly, consistent with previous observations (4, 6, 7), cysteine residues are present in regularly spaced intervals, primarily in conserved domains in all Msg proteins (Fig. 1), which likely contribute to their secondary structure. In addition, a Proline-rich motif is present in the spacer region between the M6 and C1 domains in Msg-A family and the carboxyl end of Msg-B, -D and -E families, with the average frequency of proline within this motif (20-140 aa) ranging from 18% in *P. jirovecii* Msg-A1 subfamily to 55% in *P. jirovecii* Msg-D subfamily (6-82 proline residues in each member). Furthermore, a threonine-rich motif is present in the C-terminus of Msg-A, -D and -E families (Fig. 1).

Of note, the C-terminus of Msg-D proteins has an exceptionally high content of proline, serine and threonine. One such region in protein T551_03667 includes 15 copies of a tetrapeptide repeat motif (PTDT). A similar region (rich in proline, serine and threonine) is also present in the kexin family as noted previously (1, 8) though there is otherwise no similarity between the kexin and Msg-D families.

The C-terminus of Msg-E (p55) proteins are also rich in Glu (27.2%) in addition to Pro (17.5%) and Thr (12.9%) as described above. This is consistent with previous studies of p55 in *P. carinii* and *P. murina* (9, 10), These three residues form various tandem repeat motifs present in most members of this family. For examples, three members in *P. jirovecii* (T551_01653, T551_02609 and T551_02950) each contains 13 to 31 copies of PE; another member (T551_02050) contains 6 copies of PE followed by 8 copies of PEEE. In *P. murina,* the three almost identical genes (PNEG_02059, PNEG_02419 and PNEG_03592, termed p57) each contains 8 copies of a hexapeptide repeat motif PKKED(E/D) in the center and 2 to 4 copies of PAGE repeats near 16 copies of a tetrapeptide repeat motif P(T/V/A/I)(E/K)E near the C-terminus (11). These repeats showed no intra-strain variation in the repeat copy number when aligning with Illumina reads. The homologs to these 3 genes in *P. wakefieldiae* also have these two types of repeats though with fewer copies.

Similar to the hexapeptide repeat motif in the p57 genes described above, one of the 6-gene cluster in *P. murina* (PNEG_03435), which has an extension of ~580 bp in the 3' end compared to other 5 genes, encodes 24 copies of an octapeptide repeat motif (PNDEVFGK, where underlined positions are less frequently occupied by E or S).

The roles of these enriched amino acids or motifs in *Pneumocystis* are unknown. Proline-rich regions are commonly found in proteins from different organisms and in many cases are thought to function as docking sites for signaling modules (12) or as inter-domain linker regions (13). Serine/threonine-rich regions are predicted to be *O*-glycosylated in secretory proteins encoded by diverse fungal genomes (14).

**Characterization of the mRNA expression of *P. murina msg*-A3 genes containing UCS-like sequences.**

Total RNA was extracted from *P. murina*-infected lung tissues using the RNeasy Mini Kit (Qiagen, USA), which involved a step to remove DNA with DNase I. Approximately 1 µg RNA was reversely transcribed to cDNA using SuperScript II First-Strand Synthesis System for RT-PCR (Thermo Fisher Scientific Co. USA). Primers used are listed in Table S2. The forward primers (including 4169.f10, 2833.f9, 2444.f5, 2734.f5 and 2768.f8) are specific for the UCS-like sequences in 5 *msg*-A3 genes in *P. murina*, including PNEG_02240, _00002, _03599, _03453 and _01104, respectively (Fig. 4). The reverse primer (MSG.r2b) targets a conserved region among all *msg*-A1 and *msg*-A3 genes. Reverse-transcription PCR was performed using the AccuPrime Pfx SuperMix (Thermo Fisher Scientific Co.) following the manufacturer's instructions. PCR products were purified and sequenced by Sanger sequencing.

For each of these five *msg*-A3 genes, only one sequence was obtained that contained the specific UCS-like sequence followed by a fragment specific for the corresponding *msg*-A3 gene. These findings indicate that each of these 5 genes was expressed separately and independently from the typical UCS expression site.

**Characterization of sequence variation of UCS in different *Pneumocystis* species**

For *Pneumocystis* infecting Norway rats, house mice, rabbits, dogs, and rhesus macaques, we analyzed Sanger and Illumina data from 2-8 animals per species, and observed inter- and/or intra-strain sequence variation in the UCS only for *P. carinii*, *P. wakefieldiae*, *P. macacae* and *P. oryctolagi*. Among 4 *P. macacae* isolates examined, one (H835, GenBank accession no. MN509822) showed a single SNP (non-synonymous) in exon 1 and two SNPs in the intron compared to other 3 (GenBank accession no. MN509821). Among 8 *P. wakefieldiae* isolates examined, 1 and 3 SNPs were observed in exons 1 and 2, respectively. Additionally, a stretch of 8 bp in the intron was completely different between *P. wakefieldiae* from laboratory and wild rats (Fig. S12). Among 6 *P. carinii* isolates examined, two isolates showed intra-strain variations in a 11-bp tandem repeat element in the intron; additionally one of these two isolates harbored 2 SNPs in exon 2 compared to other isolates (Fig. S13). *P. oryctolagi* isolates showed the most extensive variation, including not only many SNPs throughout but also tandem repeat variations in three regions of the intron. Tandem repeats with a 10- or 11-bp repeat unit were present with 2 to 4 repeat units per region (Fig. 5).

**Supplementary Table S1.** Nomenclature of *Pneumocystis* involved in this study.

| Mammalian common name | Mammalian scientific name | *Pneumocystis* name | Reference |
|---|---|---|---|
| Human | *Homo sapiens* | *P. jirovecii* | Frenkel (15) |
| House mouse | *Mus musculus* | *P. murina* | Keely *et al*. (16) |
| Norway or Brown rat | *Rattus norvegicus* | *P. carinii and* | Redhead *et al*. (17) |
| | | *P. wakefieldiae* | Cushion *et al*. (18) |
| European rabbit | *Oryctolagus cuniculus* | *P. oryctolagi* | Dei-Cas *et al*. (19) |
| Rhesus macaque | *Macaca mulatta* | *P. macacae* | Denis *et al*.[a] (20) |
| Dog | *Canis lupus familiaris* | *P. canis* | English *et al*. (21) |
| Polynesian rat | *Rattus exulans* | *P. exulans* | This study[b] |
| Asian house rat | *Rattus tanezumi* | *P. tanezumi* | This study[b] |
| Müller's giant Sunda rat | *Sundamys muelleri* | *P. muelleri* | This study[b] |
| Chestnut white-bellied rat | *Niviventer fulvescens* | *P. fulvescens* | This study[b] |

[a]Named as *P. carinii* f. sp. *macacae* and simplified as *P. macacae* in this study.

[b]Named upon consultation with Dr. Konstanze Bensch (http://www.mycobank.org) following the rules of the MycoBank Database for new fungal names. These names only represent *Pneumocystis* organisms from different rat species other than the Norway or Brown rat, which can be infected with *P. carinii* and *P. wakefieldiae*; further studies are needed to confirm if these organisms represent different *Pneumocystis* species.

**Supplementary Table S2.** PCR primers used in this study.

| Primer names | Targets | Species | Sequence[a] (5'→3') |
|---|---|---|---|
| WSG.f3 | CRJE | *P. wakefieldiae* | GTCCGGGGGTTGATTATTTCA |
| WSG.r5 | *msg-A1* | *P. wakefieldiae* | TTACAAGATTCCCATCATCATC |
| 3432.f1 | PNEG_03432 homolog | *P. wakefieldiae* | GATAAAGTCAATAATTTATCAAGC |
| 3438.r1 | PNEG_03438 homolog | *P. wakefieldiae* | CAAGAAGTAAAGCTGCTTTTGCT |
| RSG.f10 | CRJE | *P. carinii* | GGTTAAGAGGCAAGCA |
| RSG.r8 | *msg-A1* | *P. carinii* | CATAACAATCACTCCCAA |
| KSG.f3 | CRJE | *P. macacae* | GGGYGGCGCGGGCGGTYAAG |
| KSG.r2 | *msg-A1* | *P. macacae* | TTAAATCATAACTGAAATAACCATTC |
| OSG.f8 | *msg-A1* | *P. oryctolagi* | CCCCAGAATTTGGCGCGGGCGG |
| OSG.r11 | *msg-A1* | *P. oryctolagi* | TTACATCAACATCCAKACAAGGA |
| OSG.f3 | 5' UTR of UCS | *P. oryctolagi* | GATTTGTGCAATTATGAGAGTTGC |
| OSG.r9 | *msg-A1* | *P. oryctolagi* | CTCGCTTCTTTTGATAACATTTGT |
| 5UTR | 5' UTR of UCS | *Pneumocystis* sp | TATATTTTTCTTGATWTCCGTCT |
| CRJE.r3 | CRJE | *Pneumocystis* sp | TGYCTSCCTCTTRACCRGC |
| 4169.f10 | PNEG_02240 | *P. murina* | CATAAATGAACAAGGACATCAAGTA |
| 2833.f9 | PNEG_00002 | *P. murina* | CGTACATAAATGAACAAGACTT |
| 2444.f5 | PNEG_03599 | *P. murina* | CTCTAAAGGCATTTGGAATCC |
| 2734.f6 | PNEG_03453 | *P. murina* | ATGAAAAGTGCATCATTTA |
| 2768.f8 | PNEG_01104 | *P. murina* | GCCGCTCTAGAAAATGT |
| MSG.r2b | *msg-A1/A3*[b] | *P. murina* | ATCCTATSTTGTCCCTTTTTATA |

[a]Non-standard nucleotides (underlined) represent degenerate bases.

[b]Shared among all *msg*-A1 genes and the 5 genes of the *msg*-A3 subfamily (Fig. 4), which contain UCS-like sequences (PNEG_02240, PNEG_00002, PNEG_03599, PNEG_03453 and PNEG_01104).

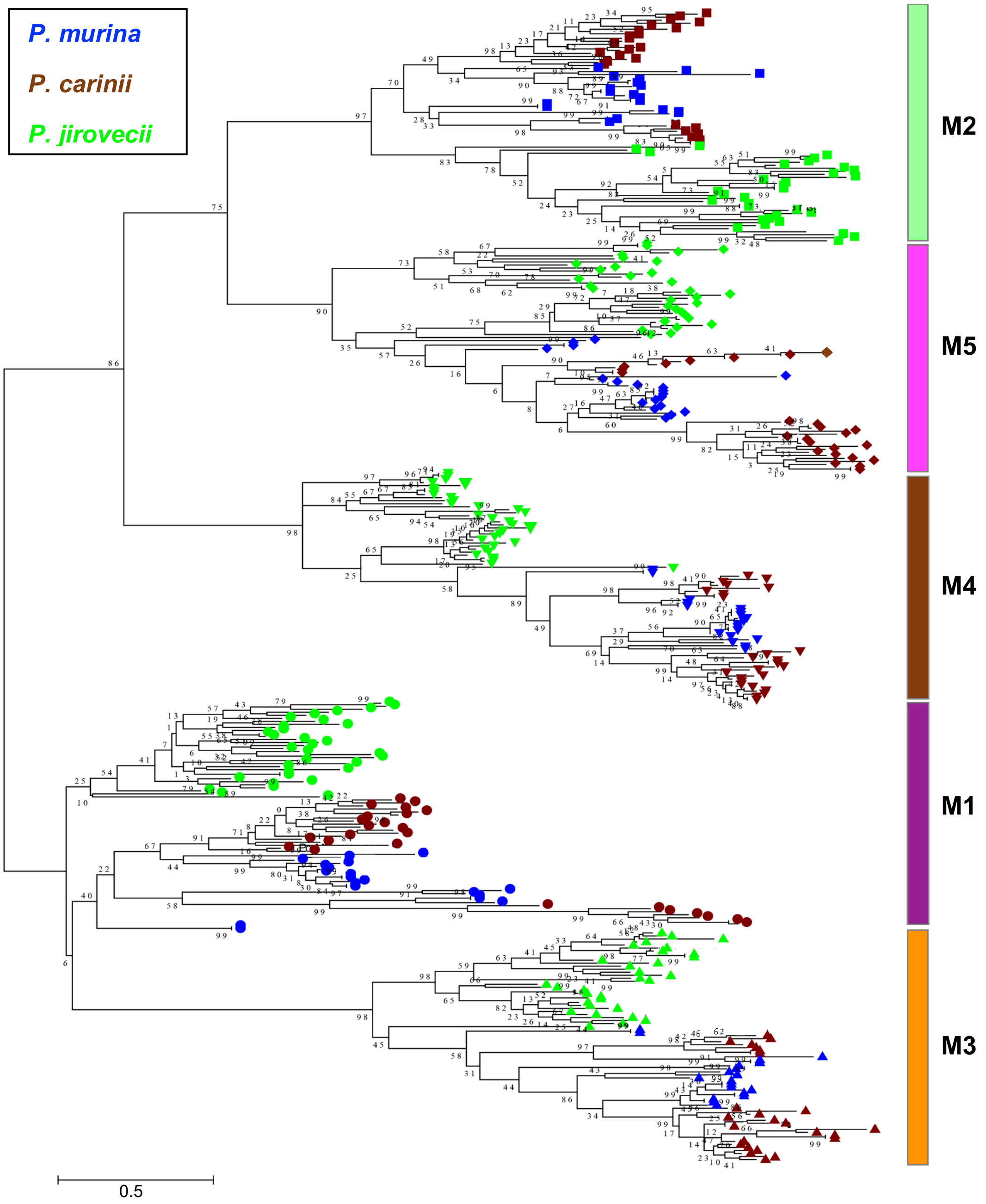


**FIG S1**. Maximum likelihood tree based on aligned but not concatenated protein sequences of Pfam MSG domains M1 to M5 from Msg proteins longer than 900 amino acids in *P. murina, P. carinii* and *P. jirovecii*. In the tree, different domains are indicated by different shapes on the right end of each branch, with different species color-coded as shown on the top left corner. The color of each domain bar is the same as in Fig. 1 in the main text. Numbers on the branch nodes indicate bootstrap support values.

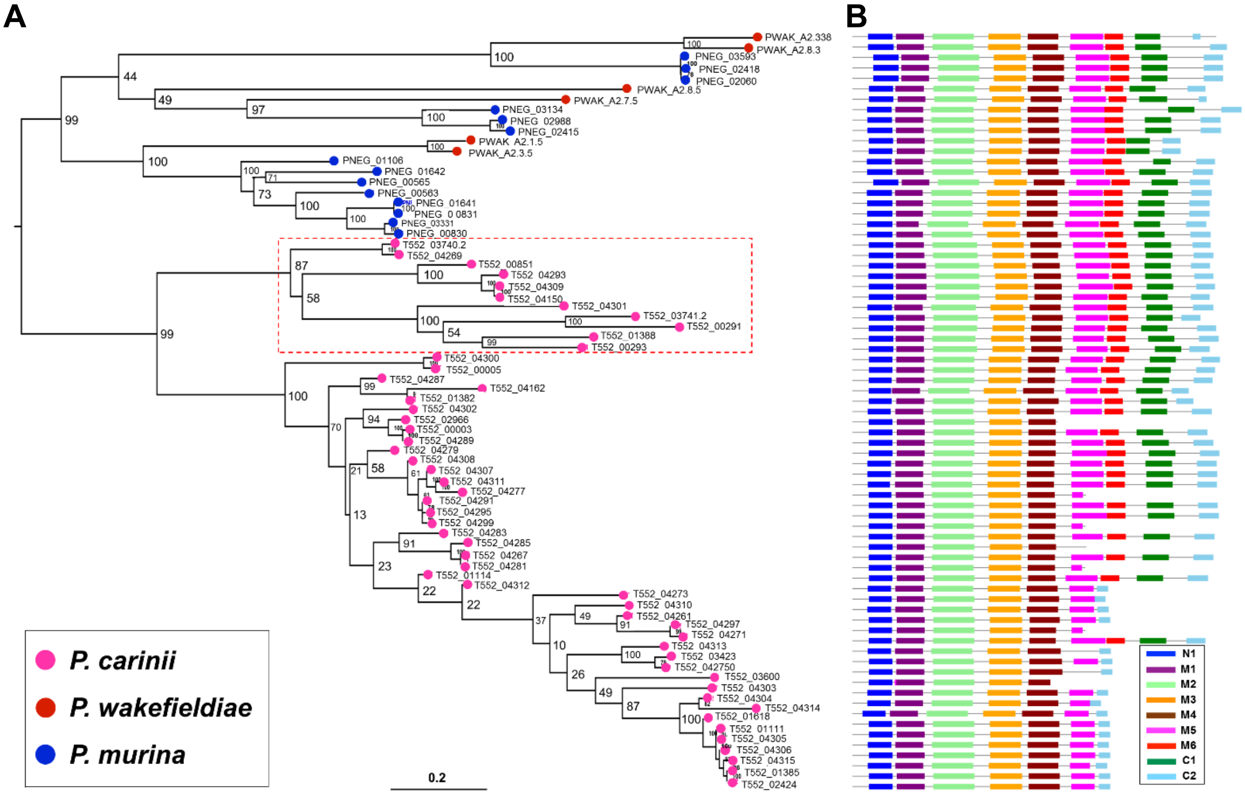


**FIG S2.** Msg-A2 (Msr) family in rodent *Pneumocystis*. **(A)** Phylogenetic relationship of the Msg-A2 family members based on full-length protein sequences. Different *Pneumocystis* species are color-coded as indicated at the bottom left corner. Numbers on the branch nodes indicate bootstrap support values. The 11 genes indicated by the red box with dashed lines form a strong clade separated from other genes in *P. carinii*, and are more closely related to msg-A1 genes as shown in Fig. S3. (**B**) Schematic representations of conserved Msg domains. Different domains are color-coded as indicated in the box at the bottom right corner. Each row corresponds to the domain structure of the corresponding protein in panel A. All sequences shown are available from Supplemental Data Set 2.

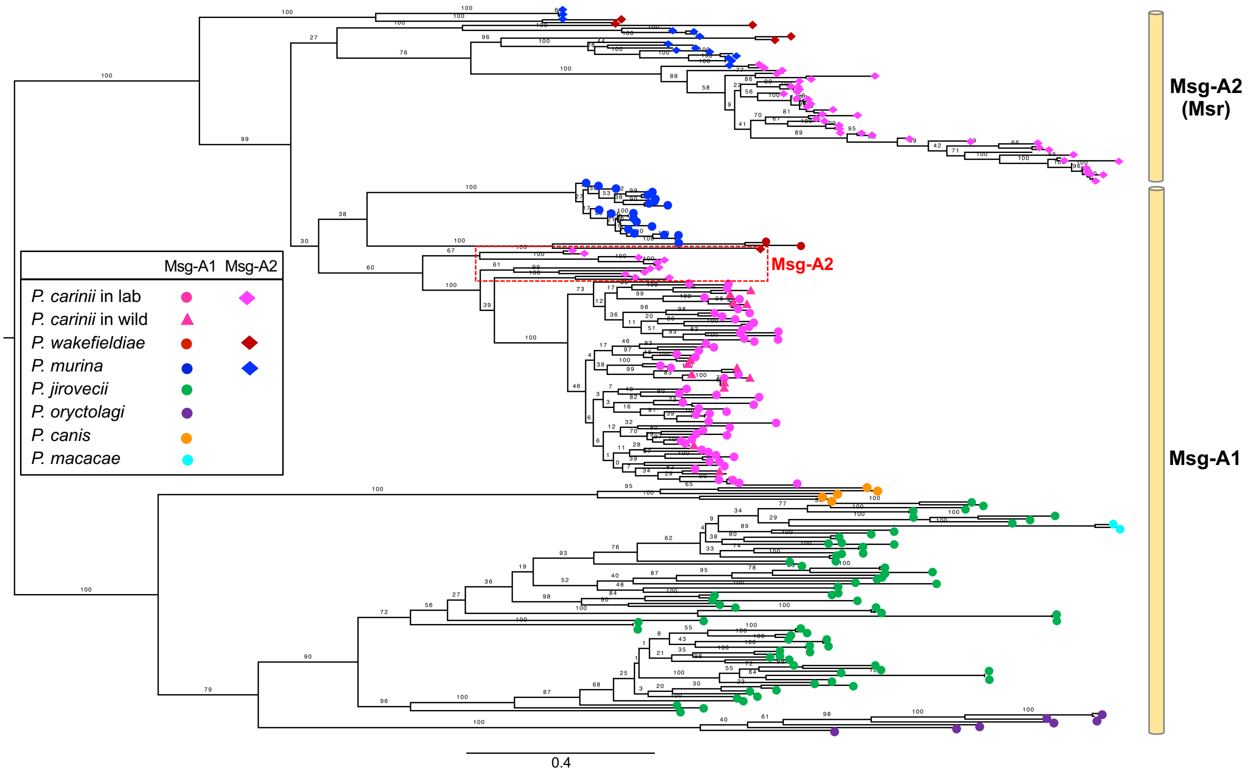


**FIG S3.** Phylogenetic analysis of Msg-A1 and Msg-A2 (Msr) subfamilies from multiple *Pneumocystis* species based on full-length protein sequences. Different *Pneumocystis* species and their Msg families are color- and shape-coded as indicated in the box within the tree. Msg-A1 genes for *P. carinii* from laboratory rats and wild rats are indicated by pink dots and pink triangles, respectively. Numbers on the branch nodes indicate bootstrap support values. All *msg*-A2 genes (only present in *P. murina*, *P. carinii* and *P. wakefieldiae*) are well separated from *msg*-A1 genes except for one in *P. wakefieldiae* and 11 in *P. carinii* from the laboratory, which are separated from other *msg*-A2 genes and clustered together within the *msg*-A1 subfamily, as indicated by the box with red dashed lines in the center.

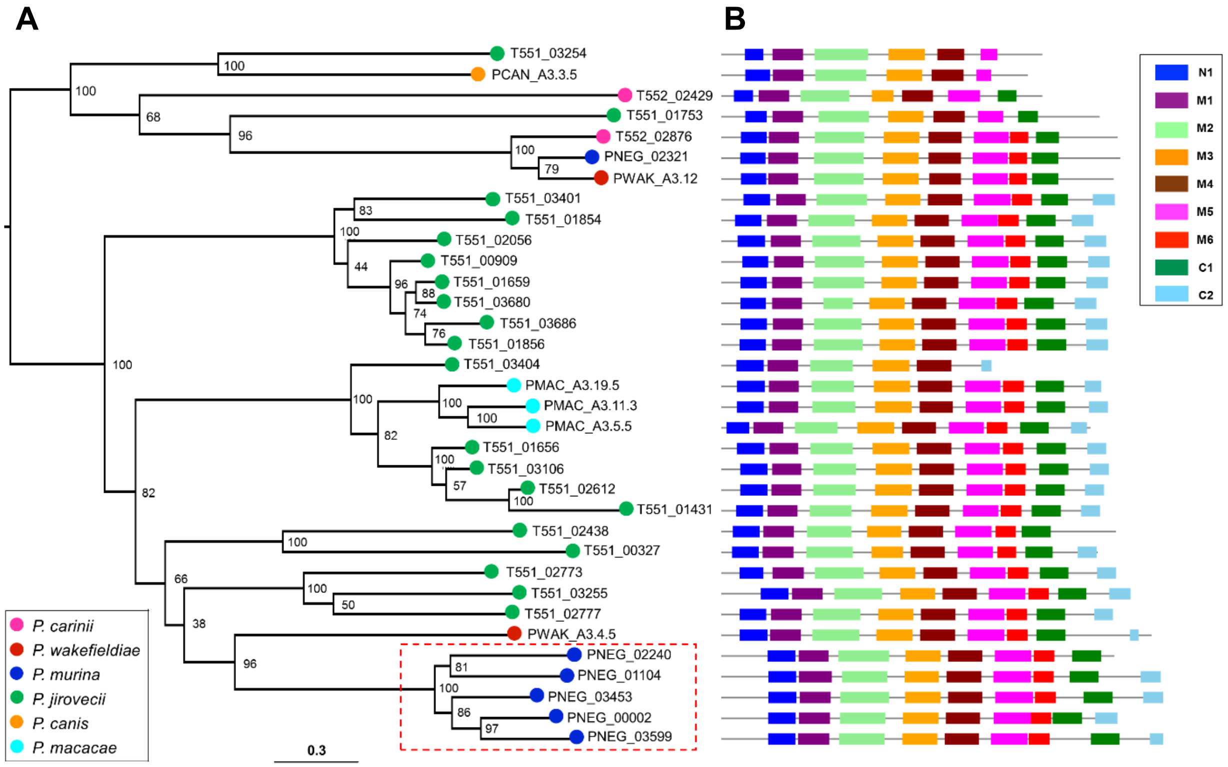


**FIG S4.** *Pneumocystis* Msg-A3 family. **(A)** Phylogenetic relationship of the Msg-A3 family members based on full-length protein sequences. Different *Pneumocystis* species are color-coded as indicated at the bottom left corner. Numbers on the branch nodes indicate bootstrap support values. The 5 genes included in the red box with dashed lines contain a UCS-like leader as shown in Fig. 4. (**B**) Schematic representations of conserved Msg domains. Different domains are color-coded as indicated at the top right corner. Each row corresponds to the domain structure of the corresponding protein in panel A. All sequences shown are available from Supplemental Data Set 3.

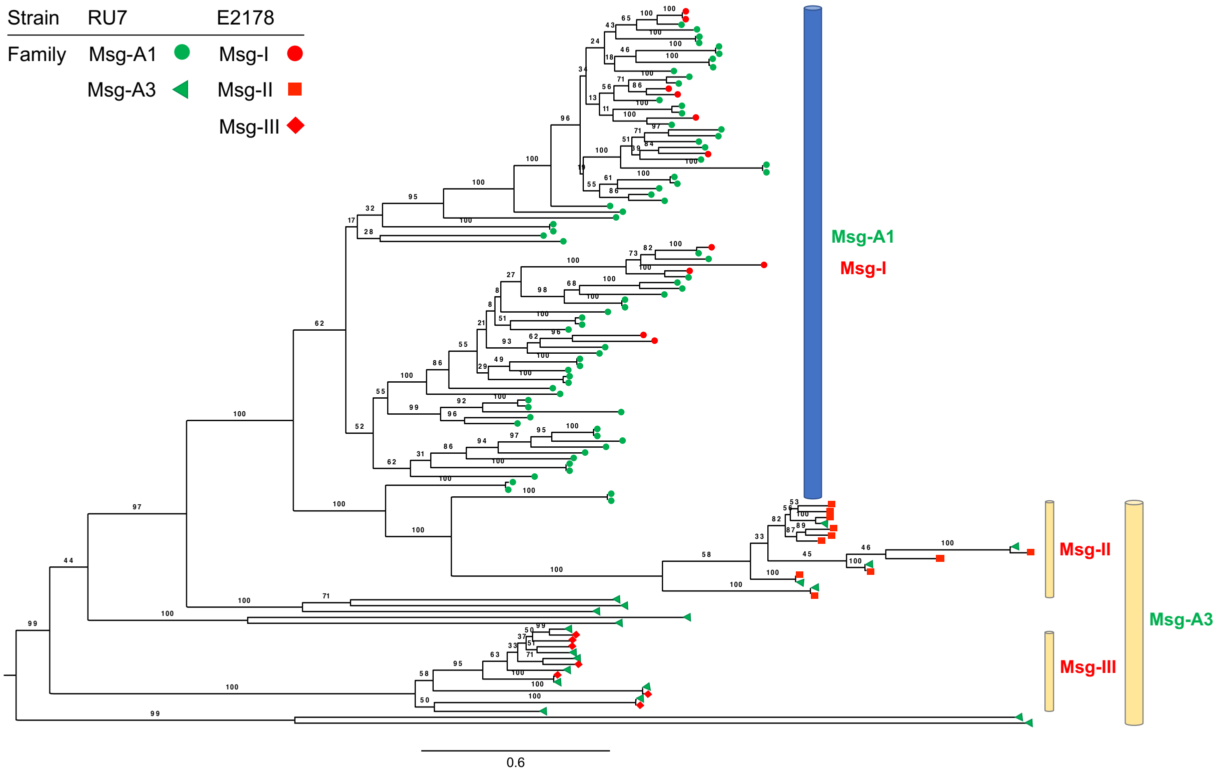


**FIG S5.** Comparison of the Msg-A1 and Msg-A3 subfamilies of *P. jirovecii* reported in different studies. Msg-A1 and Msg-A3 for *P. jirovecii* RU7 stain (indicated in green) were from our study (1) while Msg-I, -II and -III for *P. jirovecii* E2178 strain (indicated in red) were from Schmid-Siegert *et al*. (22). Different Msg families/subfamilies are indicated by different shapes and colors as shown at the top left corner. Numbers on the branch nodes indicate bootstrap support values.

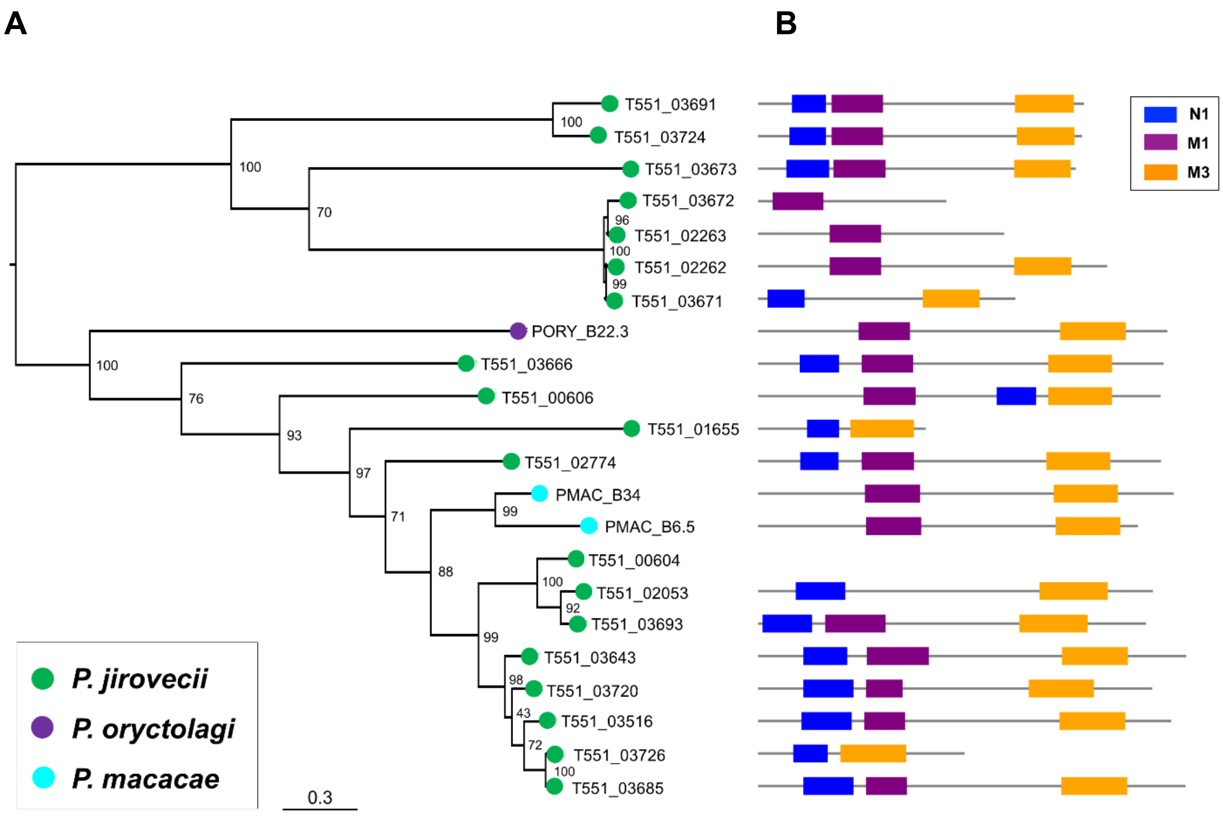


**FIG S6.** Msg-B family specific in three *Pneumocystis* species. **(A)** Phylogenetic relationship of the Msg-B family members based on full-length protein sequences. Different *Pneumocystis* species are color-coded as indicated at the bottom left corner. Numbers on the branch nodes indicate bootstrap support values. (**B**) Schematic representations of conserved Msg domains. Different domains are color-coded as indicated at the top right corner. Each row corresponds to the domain structure of the corresponding protein in panel A. All sequences shown are available from Supplemental Data Set 4.

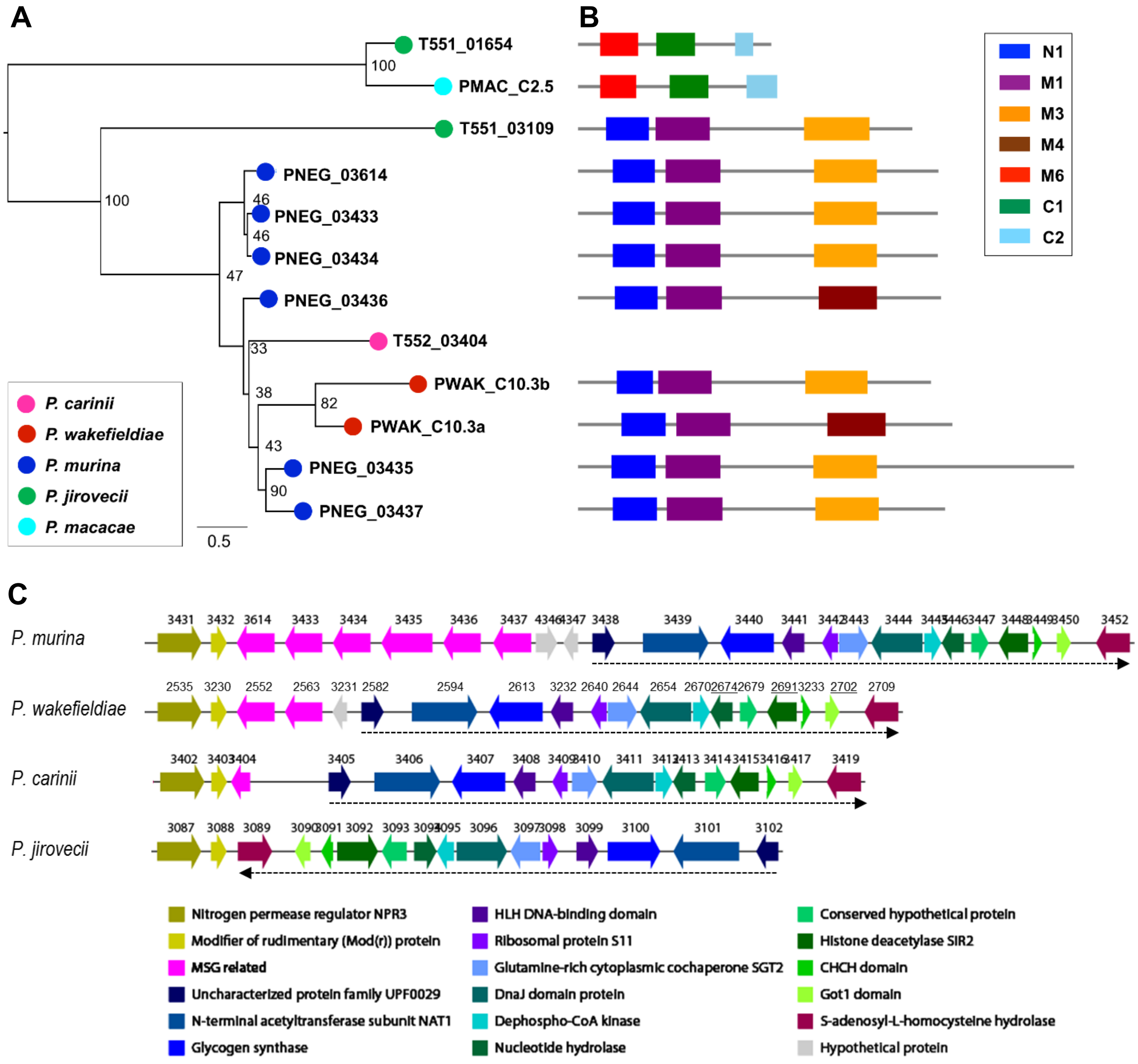


**FIG S7.** Msg-C family in five *Pneumocystis* species. **(A)** Phylogenetic relationship of the Msg-C family members based on full-length protein sequences. Different *Pneumocystis* species are color-coded as indicated at the bottom left corner in this panel. Numbers on the branch nodes indicate bootstrap support values. (**B**) Schematic representations of conserved Msg domains. Different domains are color-coded as indicated at the top right. Each row corresponds to the domain structure of the corresponding protein in panel A. All sequences shown are available from Supplemental Data Set 5. (**C**) Location of *msg*-C genes in genome assemblies (Modified from Fig. 4 in reference 1). The 6 *msg*-C genes in *P. murina* are present as a tandem cluster (shown in pink arrows) in chromosome 17, ~27kb away from the telomere (GenBank accession no. NW_017802375.1). Two copies are present in *P. wakefieldiae*. Only one partial copy of this family is present in *P. carinii* in the homologous region in scaffold 16 (GenBank accession no. NW_017264728.1) while two orthologs in *P. jirovecii* are located in two different scaffolds, with only one of them in the homologous region of scaffold 14 (GenBank accession no. NW_017264788.1). The regions flanking the 6-gene cluster of *P. murina* are conserved in all four species, except that *P. jirovecii* has an inversion (from genes T551_03089 to T551_03102) relative to the right side of the *P. murina* tandem cluster, as indicated by the dashed black line. Arrows indicate the orientations of genes, with the numbers above arrows representing the last 4 digits of the gene locus tags. The gene organization in this region in *P. macacae*, *P. canis* and *P. oryctolagi* is the same to that in *P. jirovecii* (data not shown).

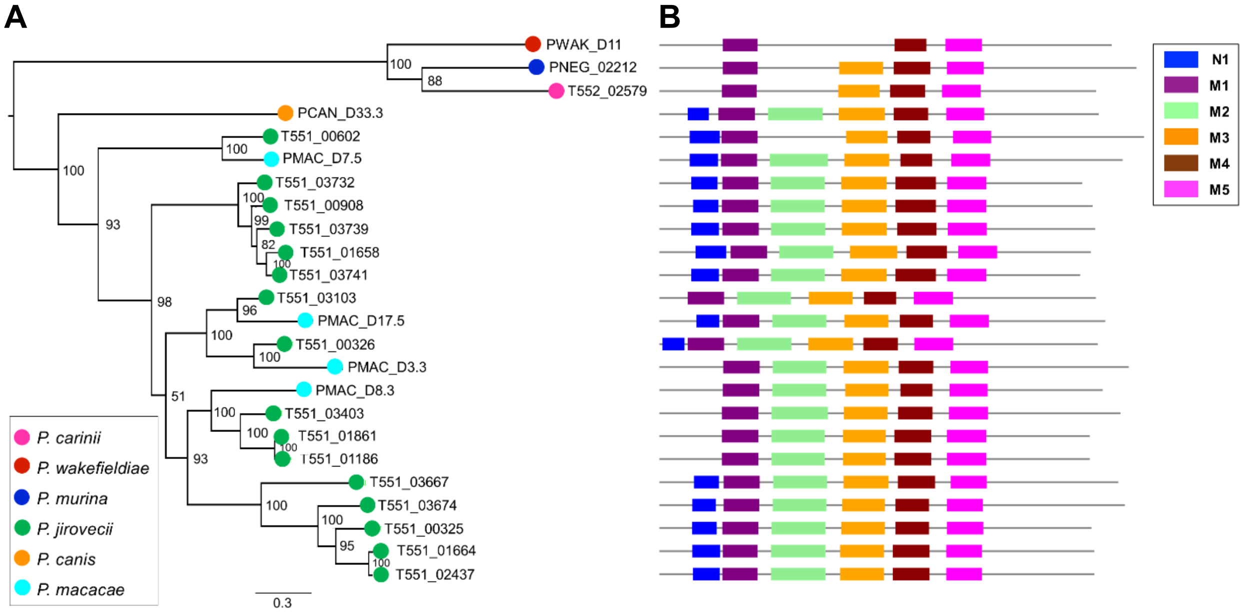


**FIG S8.** Msg-D (A12) family in *Pneumocystis* species. **(A)** Phylogenetic relationship of the Msg-D family members based on full-length protein sequences. Different *Pneumocystis* species are color-coded as indicated at the bottom left corner. Numbers on the branch nodes indicate bootstrap support values. (**B**) Schematic representations of conserved Msg domains. Different domains are color-coded as indicated at the top right. Each row corresponds to the domain structure of the corresponding protein in panel A. All sequences shown are available from Supplemental Data Set 6.

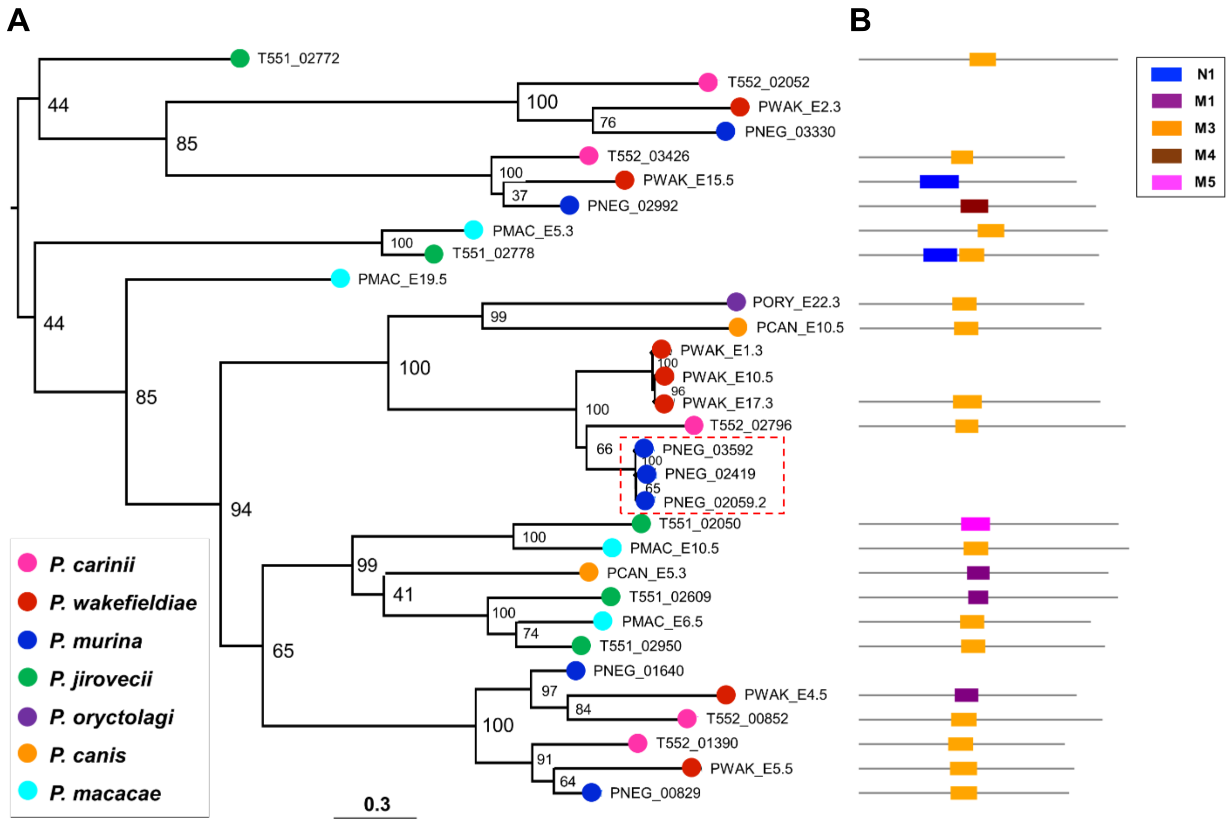


**FIG S9.** Msg-E (p55) family in three *Pneumocystis* species. (**A**) Phylogenetic relationship of the Msg-E family members based on full-length protein sequences. Different *Pneumocystis* species are color-coded as indicated at the bottom left corner. Numbers on the branch nodes indicate bootstrap support values. Three genes boxed with red dashed lines are duplicated in separate chromosomes with a nearly identical sequence and molecular size (termed p57). (**B**) Schematic representations of conserved Msg domains. Different domains are color-coded as indicated at the top right. Each row corresponds to the domain structure of the corresponding protein in panel A. All sequences shown are available from Supplemental Data Set 7.

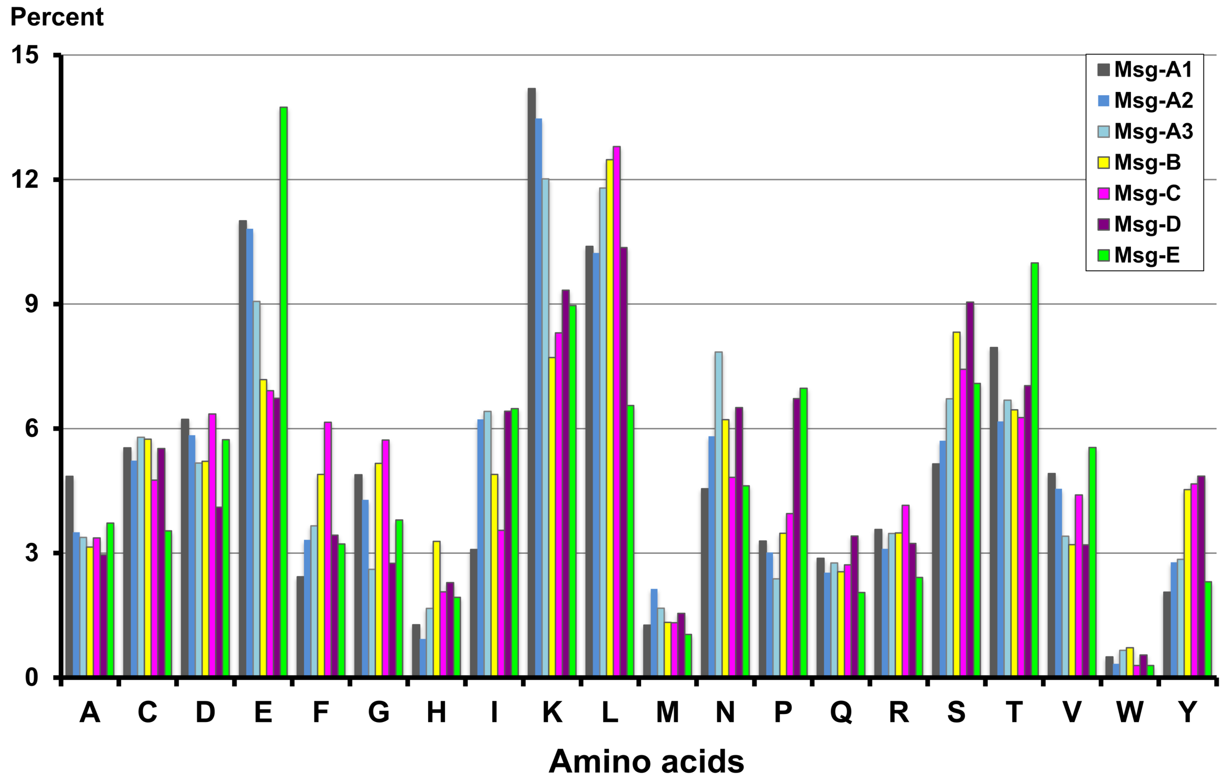


**FIG S10.** Amino acid compositions among different Msg families in *P. murina*, *P. carinii* and *P. jirovecii* with nearly fully sequenced genomes**.** Percentage was calculated using deduced amino acid sequences for all members in each family.

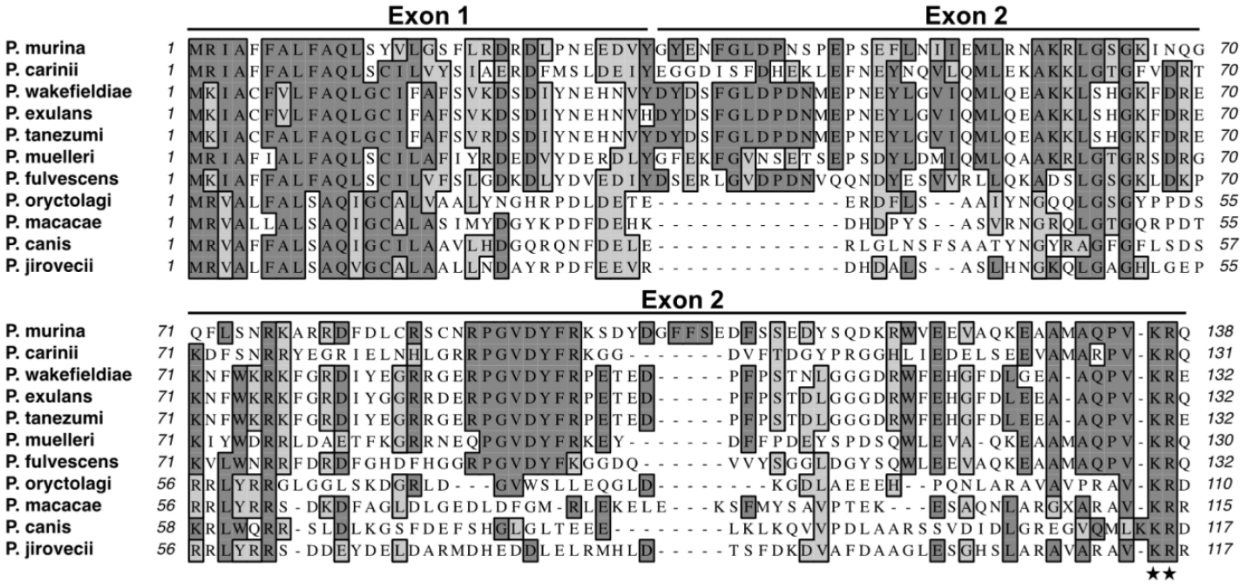

**FIG S11.** Deduced protein sequences of the expression sites or upstream conserved sequences (UCSs) of Msg-A1 in *Pneumocystis* from 11 different mammalian host species. Asterisks indicate the KR site potentially for pro-protein cleavage by endoprotease. See Fig. 3 and Table S1 for GenBank accession numbers and details about *Pneumocystis* species names.

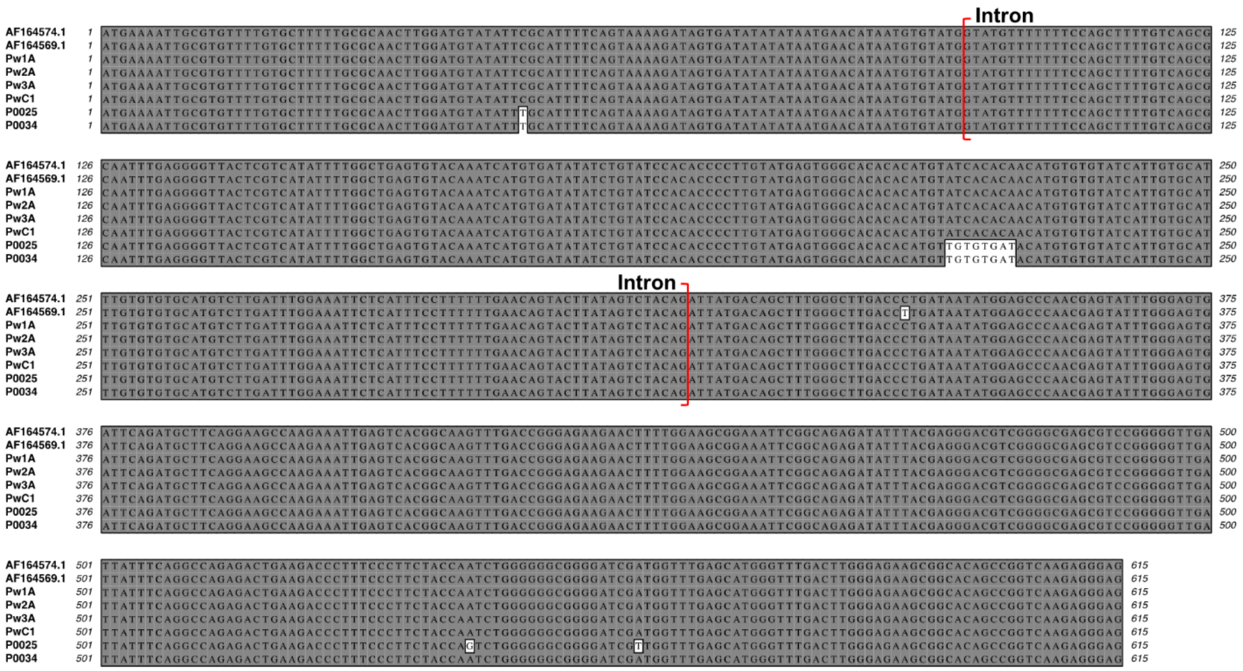

**FIG S12.** Sequence variation in the expression site (UCS) of the *msg*-A1 gene in 8 *P. wakefieldiae* isolates. Sequences for the first two isolates were reported by Schaffzin *et al*. (23), with GenBank accession nos. AF164574.1 and AF164569.1. Sequences for the other 6 isolates were obtained in this study, as determined by NGS (isolates Pw1A, Pw2A, Pw3A and PwC1 from laboratory rats) and PCR (isolates P0025 and P0034 from wild rats. GenBank accession nos. MN509829- MN509830). Numbers at either side of the alignment refer to the nucleotide positions relative to the predicted UCS translational start site. The intron is indicated in red brackets.

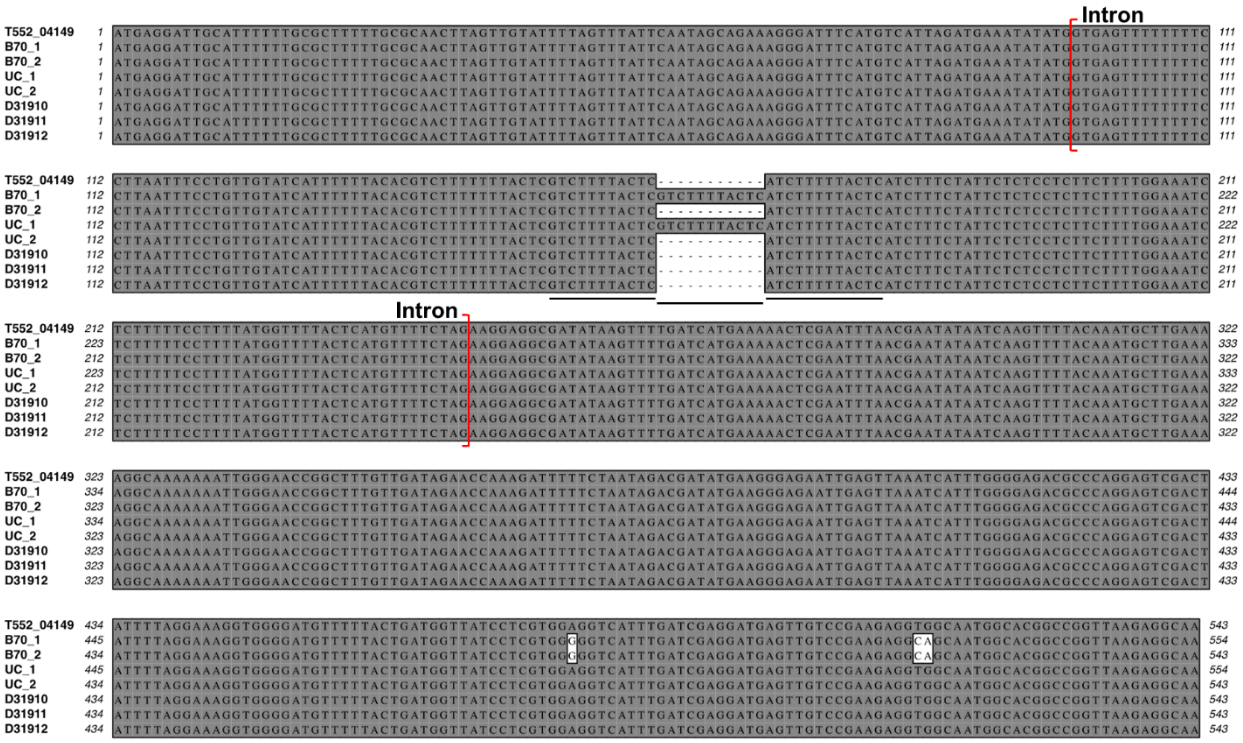


**FIG S13.** Sequence variation in the expression site (UCS) of the *msg*-A1 gene among *P. carinii* isolates. The first sequence is the *P. carinii* UCS gene from the *P. carinii* genome assembly (1). The last 3 sequences indicated by GenBank accession no. D31910-D31912 were reported by Wada *et al*. (24). Sequences B70_1 and B70_2 were assembled in this study using previous NGS data from one rat (1), available from GenBank accession nos. MN509813 and MN509814, respectively. Sequences UC_1 and UC_2 were assembled in this study using Sanger sequence reads from http://pgp.cchmc.org (25), available from GenBank accession nos. MN509815-MN509816, respectively. The 11-bp tandem repeat unit is underlined. Numbers at both sides of the alignment refer to the nucleotide positions relative to the predicted UCS translational start site.
